# SEA-Lab: Experiment-grounded recursive self-improvement in an automated laboratory

**DOI:** 10.64898/2026.11.12.766090

**Authors:** Yuliang Jiang, Ruijie Gong, He Chen, Xin Wang, Bin Zhang, Qiujun Wang, Tengfei Zhang, Mengya Fan, Xueya Mao, Zhenling Hou, Yang Hu, Yuan Gao, Cheng-Zhi Wang

## Abstract

AI scientists can generate hypotheses, plans, and protocols. Translating these outputs into reliable experiments requires adapting them to local equipment and resources using feedback from observed outcomes. A key challenge is to retain laboratory execution feedback as reusable knowledge for subsequent planning. We present SEA-Lab (**S**elf-**E**volving **A**utonomous Lab), which combines an environment-grounded Dry-Lab Harness that generates protocols and machine-readable procedures, a wet-lab execution harness that coordinates heterogeneous devices and records execution traces and experimental measurements, and a recursive self-improvement mechanism that uses physical feedback to revise reusable execution policies and experimental strategies. Across three rounds of review on 219 tasks, protocol-review pass rates reached 90.41%. On 66 automation-eligible tasks, mean virtual-validation pass rates increased from 16.7% to 87.7% after two revisions. On two liquid handlers, runtime fell by 45.8% for plate replication, 34.1% for well rearrangement, and up to 68.6% for sample normalization. We demonstrated the system in a phage-display experiment linking four automated experimental steps: molecular cloning, phage titration, sample normalization, and ELISA screening. Of 96 starting samples, 93 yielded a sequence-correct clone, and primary ELISA screening identified ten candidates. These results illustrate how laboratory feedback can guide planning around local resources, refine execution on different laboratory instruments, and inform method revision based on experimental results.

## 1 Introduction

Large language model (LLM)-based AI scientists are beginning to integrate literature retrieval, hypothesis generation, experimental design, and data analysis into a single research loop [1]. Systems such as Robin [2] and Co-Scientist [3] can assemble literature evidence, propose hypotheses and research plans, and use experimental validation or result analysis to support subsequent decisions. Other platforms extend this process to physical experiments by connecting agentic planning and protocol translation to robotic execution [4, 5]. Wet-lab feedback also informs protocol revision in ProtoPilot [6], while iCloud-Biofoundry [7] connects natural-language research goals to remote biofoundries and returns measurements to inform subsequent designs. A central challenge is to adapt experimental plans to local laboratory conditions and retain feedback from execution and experimental outcomes as reusable knowledge for subsequent planning. We frame this problem as three related mismatches.

First, a **knowledge–environment mismatch** exists between general experimental knowledge and local conditions. The literature and standard protocols describe the principles and main steps of a method, but they may not fully specify how to implement it with a particular combination of instruments, consumables, and reagents. The tacit operational knowledge in a manual protocol must be made explicit when the protocol is transferred to an automated platform, and adapted to device specifications and experimental goals [8]. The same objective can be met by several valid procedures, but their feasibility in a given laboratory is constrained by device capabilities, consumable compatibility, reagent availability, and operating limits. A protocol suited to a particular laboratory must specify which local instruments, consumables, and reagents to use, and how to work within their operating limits [9].

Second, a **plan–execution mismatch** arises because a locally feasible protocol may behave differently during physical execution. Planning relies on estimates of device state, operation duration, action reliability, and resource occupancy. These estimates may not match actual execution. Roboticarm travel, plate-transfer times, positioning offsets, and inter-device waits can affect workflow completion and throughput [10]. Platform-specific script generation, pre-execution validation, and runtime control improve the feasibility of commands [11]. Feedback from physical execution is needed to update the system’s model of device capabilities, operation durations, and workflow handoffs.

Third, a **protocol–outcome mismatch** arises because faithful execution of a valid protocol does not guarantee a biologically successful or interpretable result. Outcomes must therefore be assessed using biochemical or functional readouts appropriate to the research objective [12, 13]. Reagent variation, sample physiological state, upstream processing, and the experimental environment can all produce substantial differences in performance. An uninformative or technically inadequate result may arise from a specific operation, a parameter setting, workflow design, or the broader experimental strategy. The AI scientist must assess which level, if any, warrants revision.

To address these mismatches, we developed SEA-Lab (**S**elf-**E**volving **A**utonomous Lab), which combines an environment-grounded Dry-Lab Harness, a Wet-Lab Harness for heterogeneous device orchestration, and a recursive self-improvement (RSI) mechanism. The RSI mechanism is designed to use execution records and experimental outcomes to revise reusable execution policies and experimental strategies for subsequent planning.

We organized the evaluation around the three mismatches: locally grounded protocol generation and virtual validation for the knowledge–environment mismatch; instrument-based liquid-handling refinement for the plan–execution mismatch; and biological method development and a phage-display workflow for the protocol–outcome mismatch.

## 2 Methods

### 2.1 System overview

SEA-Lab (Fig. 1) comprises a Dry-Lab Harness for locally grounded planning, a Wet-Lab Harness for device orchestration, and an RSI mechanism for feedback-guided revision. The Dry-Lab Harness generates experimental protocols and machine-readable procedures encoded in JavaScript Object Notation (JSON-procedures). The Wet-Lab Harness executes these procedures and records device activity and experimental outputs. The RSI mechanism is designed to use these records to revise execution policies and experimental strategies for subsequent planning.

**Fig. 1.**
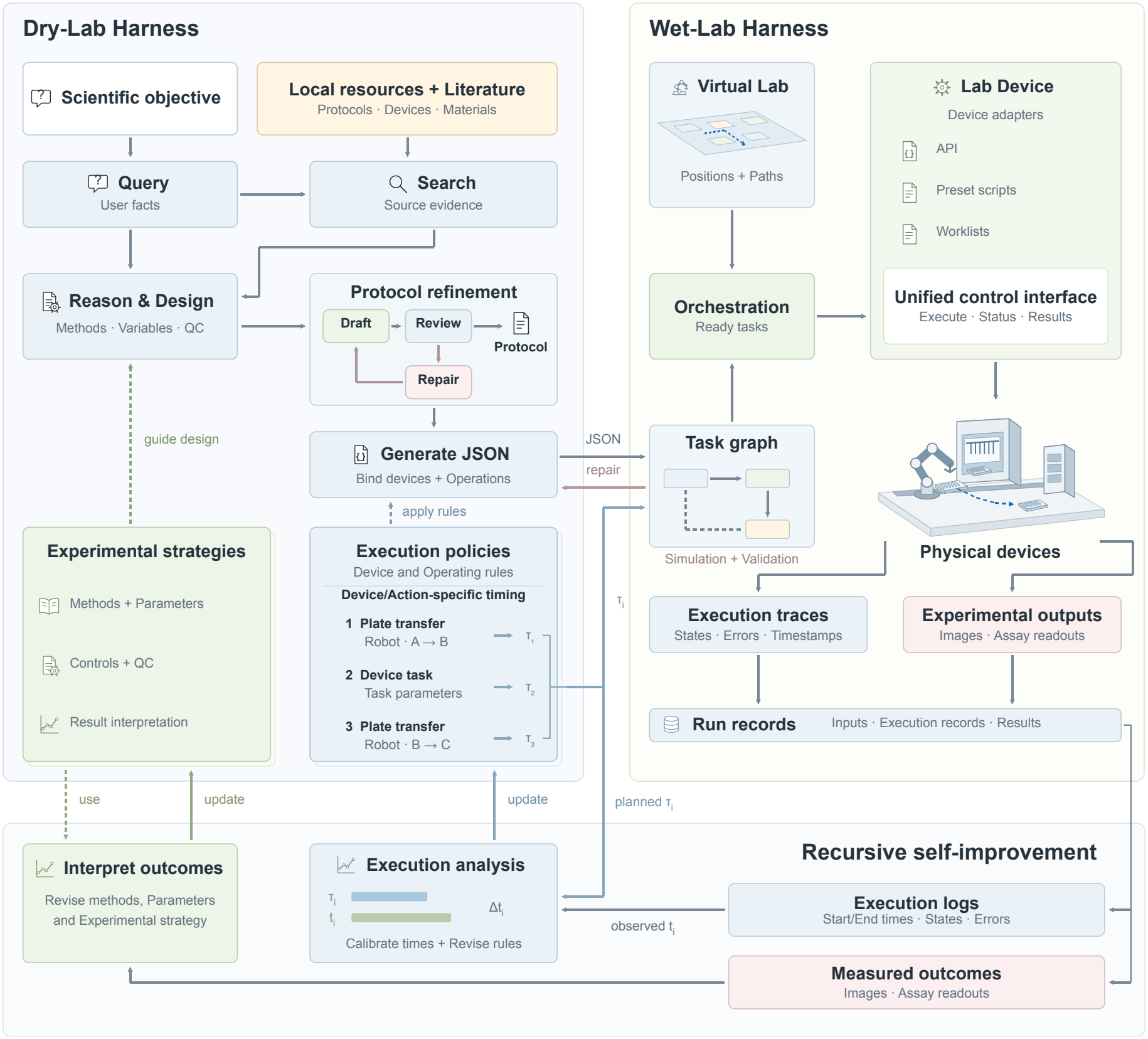
SEA-Lab architecture and feedback-guided revision of policies and strategies. The Dry-Lab Harness combines the scientific objective, *user facts*, local resources, and retrieved evidence to design and refine a protocol, then generates a JSON-procedure. Experimental strategies inform experimental design, while execution policies provide operating rules and timing estimates. The Wet-Lab Harness validates the procedure and organizes a task graph, using *Virtual Lab* positions and paths and a unified *Lab Device* interface to coordinate physical devices. Run records capture input parameters, execution logs, and experimental outputs. Execution analysis compares estimated durations *τ*_*i*_ with observed durations *t*_*i*_ to inform timing and rule revisions; outcome interpretation informs revision of experimental strategies. The feedback paths show how revisions can inform subsequent planning. Solid lines indicate processes, records, or updates, and dashed lines indicate policy or strategy use.

### 2.2 Dry-Lab Harness for grounded planning

The Dry-Lab Harness integrates a local protocol library, device capabilities and operating limits, and consumable compatibility information to ground experimental planning in the target laboratory (Fig. 1).

The *Query* module clarifies the experimental objective, available devices and materials, intended methods, and plans for preliminary experiments. Confirmed conditions are recorded as user-facts for subsequent design and procedure generation. A retrieval module (*Search*) draws on local resources and external literature to identify candidate methods, reference steps, and supporting evidence, retaining the relevant content and its sources as source-evidence. A design module (*Reason & Design*) combines these inputs with local resource constraints to select methods and define key variables, controls, and quality-control requirements.

Protocol generation produces an initial draft, which is checked and revised through *Review* and *Repair* for experimental logic, completeness, and consistency with local resources.

For the Tecan Freedom EVO 150, a Worklist Generator translates structured operations into advanced worklists (supporting script and device commands) using local execution configurations supplied by the Wet-Lab Harness (Supplementary Fig. S1 and Supplementary Section S1). It selects a compatible predefined deck layout and uses registered labware mappings and liquid classes (sets of pipetting parameters such as aspiration and dispense speeds) to generate device-specific instructions. The JSON-procedure specifies cross-device dependencies and references the generated worklists. Worklists are validated together with the main procedure for consistency with the selected layout, labware bindings, and inventory.

The reviewed protocol is translated into a JSON-procedure using confirmed user facts, registered device actions and parameter constraints, labware definitions, initial positions and inventory, and the input schema. The conversion interface requires unsupported operations and missing information to be reported explicitly, without inventing capabilities or substituting placeholder tasks for required operations (Supplementary Section S2). The resulting procedure is passed to the Wet-Lab Harness for validation and execution.

Experimental strategies guide *Reason & Design*, while execution policies guide protocol-to-JSON conversion; their contents and feedback-guided revision are described in Section 2.4.

### 2.3 Wet-Lab Harness for device orchestration and observation

The Wet-Lab Harness is the execution and observation layer of SEA-Lab (Supplementary Fig. S2, Supplementary Section S3, and Supplementary Table S3). It receives a JSON-procedure and organizes experimental steps into tasks with dependencies. *Lab Device* provides device invocation and state queries, and *Virtual Lab* provides spatial positions and transfer configuration.

#### Heterogeneous device abstraction and capability boundaries

Because devices differ in their operating logic, *Lab Device* provides a unified invocation layer above each device’s native control interface so that instruments with different control modes can be connected. This interface exposes common command invocation, device-status queries, and result reporting across instruments. The device-specific implementation translates these calls into the control mode supported by each instrument: an application programming interface (API), a preconfigured script, or an advanced worklist.

Devices are grouped by their role in the workflow. Plate-handling devices, including robotic arms and linear rails, transfer and position plates; experimental devices, including liquid handlers, automated incubators, and refrigerated high-speed centrifuges, perform pipetting, cultivation, and other operations. The scope of device invocation is bounded by existing control capability: for API-controlled operations (e.g., robotic arms), executable tasks are limited to the settings exposed by the API. For script-controlled operations (e.g., the thermal cycler), executable tasks are limited to existing scripts. For worklist-controlled operations (e.g., liquid handlers), the Dry-Lab Harness generates a worklist at the configured storage location and supplies its path for execution. Device positions, available plate slots, and other conditions that must be configured in advance are set before execution, and experimental tasks are executed within the range permitted by the local platform.

#### Experimental layout and position initialization

The physical platform layout is shown in Supplementary Fig. S3. *Virtual Lab* provides device layouts, logical plate slots, access permissions, and transfer points (Supplementary Section S3.2 and Supplementary Table S4). These definitions map device and labware identifiers in a JSON-procedure to physical locations. Layout definitions are maintained separately from labware-instance state, which records identity, location, labware specification, lid status, and occupancy. The relevant layout and initial labware state are configured before execution. Configured slot capacity does not imply that labware is present.

#### Structured task parsing and workflow orchestration

The Wet-Lab Harness parses each JSON-procedure into device-operation, labware-transfer, and wait or software-processing tasks (Supplementary Section S3.3 and Supplementary Table S5). Dependencies declared through depends on define a directed acyclic task graph; task order in the JSON array does not determine execution order. A task becomes eligible to start only when its predecessors have succeeded, the required labware is in place, the device is ready, and the necessary resources are available. Explicit dependencies remain binding when additional condition-based dependencies are introduced.

#### Execution validation and runtime feedback

The Wet-Lab Harness performs checks at submission, scheduling, execution, and completion (Supplementary Section S3.4 and Supplementary Table S6). It separately reports schema validity, pre-execution validation, virtual-execution status, and acceptance against independently specified objectives. Additional checks depend on explicitly supplied models or constraints; missing declarations are not treated as evidence that a requirement has been satisfied. ERROR diagnostics block validation, whereas WARNING diagnostics are permitted in normal mode but block validation in strict mode. Structured diagnostics identify the rule, severity, stage, and affected task or field. Failed candidates are revised using these diagnostics and revalidated while retaining the original experimental objective and confirmed inputs.

#### Completion evidence and run records

Command acceptance is distinguished from action completion. The orchestration layer queries execution status using the corresponding execution identifier and verifies completion evidence. For data-producing operations, returned records must also match the current execution. Downstream tasks cannot proceed without confirmed completion of their prerequisites. Confirmed actions update labware locations, resource occupancy, and material state. Procedure and configuration snapshots, execution records, and state history are retained for traceability, visualization, and replay (Supplementary Fig. S2 and Supplementary Section S3.5). Interrupted runs retain confirmed state changes and mark unresolved states as uncertain; potentially completed actions are not automatically replayed while their execution state remains uncertain.

#### Human participation

The researchers provide the experimental objective and clarify task requirements. Before a physical run, they place consumables in the designated positions and check that the plate layout on the platform matches the configuration used by the Wet-Lab Harness. They also participate in reviewing generated protocols and candidate execution policies proposed by the RSI mechanism.

### 2.4 Feedback-guided revision of policies and strategies

The RSI mechanism is designed to retain execution policies and experimental strategies for subsequent planning and revise them as further observations become available. This differs from feedback-guided procedure refinement, in which a candidate JSON-procedure is repaired within its current task.

SEA-Lab uses three sources of feedback. Protocol-review feedback identifies deficiencies in experimental rationale, completeness, and compliance with local resources. Virtual-validation feedback identifies violations of device, position, dependency, timing, and analysis constraints in a JSON-procedure. Physical feedback comes from device states, time-stamped execution logs, and experimental measurements captured in run records. These sources differ in what they establish: review assesses the proposed plan, virtual validation assesses compliance with encoded constraints, and physical observations describe actual execution and experimental outcomes. Physical feedback, including errors and interruptions, is retained in the run record and linked to the corresponding execution identifier. Labware locations and material states are updated only after the relevant action or step is confirmed complete, as described in Section 2.3

Run-record analysis follows two complementary paths. Execution analysis uses device states, errors, and operation logs to identify problems in device operation and workflow handoffs. Comparisons between simulated and observed durations inform revisions to timing estimates and representations of the local execution environment. Outcome analysis examines experimental measurements to guide changes in method selection, experimental conditions, and interpretation of results. These analyses inform revisions to execution policies and experimental strategies, respectively.

In the RSI design, adopted execution policies guide subsequent procedure generation, while experimental strategies guide subsequent experimental design. New runs provide further feedback for revision. The evaluation below distinguishes repair within a task from reuse between rounds by tracking whether an adopted pipetting policy becomes the reference for subsequent comparisons.

### 2.5 Experimental design and evaluation

We evaluated SEA-Lab in five areas: locally grounded protocol planning, procedure validation, physical workflow execution, feedback-guided refinement, and end-to-end biological application. We report virtual-validation results, instrument performance, and biological outcomes separately to distinguish compliance with encoded execution constraints, completion of automated workflows, and achievement of experimental objectives.

#### Localized protocol generation

We constructed a protocol-generation benchmark comprising 219 tasks assigned to eight mutually exclusive primary experimental domains (Supplementary Fig. S4A). The researchers prepared and reviewed reference protocols according to local devices, reagents, and consumables, and converted them into experimental intents. A representative experimental intent is shown in Supplementary Fig. S4B. A *Query* agent then submitted them item by item to the system. Three rounds of generation and evaluation were performed on the same set of tasks. Error analyses from the *Review* agent in each round informed protocol generation in the next round.

Generated protocols were scored independently by human scientists and a *Review* agent using a common rubric covering five dimensions: requirement consistency, scientific rationality, completeness and executability, quality control and exception handling, and clarity and traceability. Requirement consistency included compliance with confirmed conditions and local resource constraints. Human scores were weighted two-thirds and agent scores one-third. A protocol passed only when both parties judged it acceptable and the composite score was at least 80 (Supplementary Section S4 and Supplementary Table S7). Metrics were the pass rate, composite score, dimension scores normalized to their maximum, and the number of protocols with critical issues.

#### Protocol-to-JSON conversion and validation

We repeated the evaluation seven times on the same set of 66 automation-eligible tasks selected from the protocol-generation benchmark. In each evaluation, the Dry-Lab Harness generated an initial JSON-procedure for each task (R0), and the Wet-Lab Harness checked it through simulated execution and software rule checking in *Virtual Lab*. The system used validation feedback to revise procedures that failed, with subsequent revisions denoted R1, R2 and so on. We recorded the validation pass rate at each round, the number of revisions required for first pass, and the number of diagnostic issues by category. We compared R0 and R2 for each task within the same evaluation, recording which initially failing tasks passed and whether any initially passing tasks failed. We also compared the number of revision rounds for preparation workflows and for tasks involving quantitative analysis. Diagnostics were grouped into eight categories (Supplementary Fig. S5, Supplementary Section S3.4, and Supplementary Table S6). Within each evaluation and revision round, each procedure was counted at most once per category. Categories were non-exclusive, whereas the total number of failed procedures was deduplicated across categories. Unmeasured rounds were distinguished from observed zero counts. This evaluation reflects only executability under the defined simulation environment and software rules.

#### Liquid-handling policy and scheduling refinement

For pipetting tasks on a liquid handler, candidate policies varied tip handling, aspiration grouping, flow rate, blow-out, and transfer order within task-specific constraints. For each experiment performed on the Opentrons OT-2, the agent could propose up to 10 candidate policies. Each candidate was tested at least three times against the current policy, with runtime as the only performance metric compared. A paired comparison counted as an improvement only if the candidate reduced runtime by more than 10% relative to the current policy. For well rearrangement on the Tecan Freedom EVO 150, the source–destination well mappings, the liquid class, and transfer volume were held constant within each layout, while source-well visit order, channel assignment, destination dispense route, and tip usage were allowed to vary. A simulator was built from real run logs to support generation and revision of candidate schedules. The built-in strategy for the basic layout served as the initial reference, and candidate schedules were examined in the basic layout and four additional sourceplate layouts. The well-rearrangement runtime comparisons use recorded instrument runs. For phage titer normalization, we compared five scheduling versions (v0–v4) across three initial layouts, labeled Original, Layout 1, and Layout 2, with one instrument run per layout–version combination. We compared summed action times, aspiration counts, and tip use from the available summaries. Summed action time is the sum of the reported action categories and is distinct from the total-runtime field in these summaries; missing component entries were not interpreted as zero-duration measurements.

#### Experimental method development

We examined scientific decision-making through iterative *ccdB* negative-selection vector design and phage titer assay development. Vector design used recovery of intact plasmids and background colony formation relative to matched positive controls to guide subsequent regulatory-element combinations. Titer-assay development used colony growth and the suitability of colony counts for titer estimation to guide changes in dilution and mixing, reagents, and host-cell state. These cases describe how experimental observations informed subsequent choices.

#### End-to-end phage-display experiment

The batch experiment linked molecular cloning, phage titration, titer-guided normalization, and ELISA screening. Molecular cloning used 96 samples; we recorded colony recovery rate, the proportion of colony-positive samples with at least one correct clone, and construction success rate over all starting samples. Up to two colonies per colony-positive construct were submitted for Sanger sequencing. A construct was considered successful when at least one clone matched the expected sequence. Titer measurement included 95 samples: the 93 sequence-correct clones, a positive control, and a negative control.

ELISA screening included 93 clones, each measured once on each of two plates containing positive, negative, and blank controls. Inter-plate association was assessed using Pearson and Spearman correlations across clone measurements, excluding control wells. Signal-to-negative-control ratios (*S/N* ) were calculated separately for each plate as the sample OD_450_ divided by the negative-control OD_450_ in G12, without blank subtraction. Clones with *S/N >* 3 on both plates were designated as primary-screen candidates and ranked by mean OD_450_. Experimental procedures and titer normalization are described in Section 2.6.

#### Workflow execution on heterogeneous devices

The cloning, titration, and ELISA workflows were used to evaluate handoffs among plate transfer, pipetting, cultivation, plate washing, and plate reading. Titer-guided normalization was part of sample preparation for ELISA. The system recorded step start and end times, device status, interruptions, and completion. For each of the three workflows, the completion rate was calculated as completed runs divided by recorded runs. Workflow durations were also reported. Logs and task timelines were used to analyze problems in device invocation, spatial positioning, and cross-device handoffs.

### 2.6 Wet-lab workflows

The biological workflow proceeds from molecular cloning to phage infection titer measurement and titer-guided ELISA screening (Supplementary Figs. S6 and S7; Supplementary Section S5). The Wet-Lab Harness coordinates the labware carousel, liquid handler, thermal cycler, incubator, colony picker, microplate washer, and microplate reader according to task dependencies. The following procedures describe vector preparation and the experimental steps; evaluation criteria are defined in Section 2.5.

#### Vector construction and cloning

The pDWPD V1–pDWPD V8 series varied the promoter and ribosome-binding site (RBS) controlling *ccdB*. Construction of the variants was attempted by an external supplier in the *ccdB* -tolerant host *E. coli* DB3.1, and Sanger sequencing was used to check the promoter and regions adjacent to *ccdB*. Recovery of an intact expected plasmid was the criterion for successful construction. The recovered variants pDWPD V1, pDWPD V2, and pDWPD V8 were chemically transformed into *E. coli* TG1 to assess negative selection, using 2 ng and 20 ng plasmid inputs and matched positive controls in two independent experiments. Cells were recovered in 200 *µ*L LB for 1 h and plated on ampicillin-containing medium, followed by incubation at 37 ^*?*^C for 16 h. Background colony rate was calculated relative to the matched positive control.

For cloning with pDWPD V8, NheI and NotI digestion removed *ccdB*, and an insert prepared by primer annealing was ligated using T4 DNA ligase in a 10 *µ*L reaction at 25 ^*?*^C for 1 h. The product was transformed into TG1 using the recovery and cultivation conditions above. Clones recovered in the batch application were submitted for Sanger sequencing as described in Section 2.5.

#### Phage titer measurement and ELISA

Phage infection of TG1 was quantified by counting resistant colonies after spot plating. Assay development compared changes in dilution and mixing, reagents, and host-cell state, including the use of TG1 selected on M9 medium. Before ELISA, samples above 4 *×*10^9^ infectious units per milliliter (IU/mL) were diluted toward that target, whereas samples at or below it were used undiluted. Pre-blocked phage preparations were incubated in target-protein-coated, blocked wells, followed by washing and detection-antibody incubation. After a final wash, substrate was added, and the reaction was stopped before absorbance measurement at 450 nm (Supplementary Figs. S6C and S7H). Each of the 93 clones was assayed on two plates, each containing positive, negative, and blank controls.

## 3 Results

### 3.1 Grounded planning aligns experimental knowledge with local laboratory resources

#### 3.1.1 Dry-Lab Harness generates protocols grounded in local devices and consumables

To assess the knowledge–environment mismatch, we first evaluated localized protocol generation on the 219-task benchmark (Fig. 2A–C, Supplementary Section S4, and Supplementary Table S7). Three rounds were evaluated on the same tasks, with *Review* agent error analyses informing protocol generation in the next round.

**Fig. 2.**
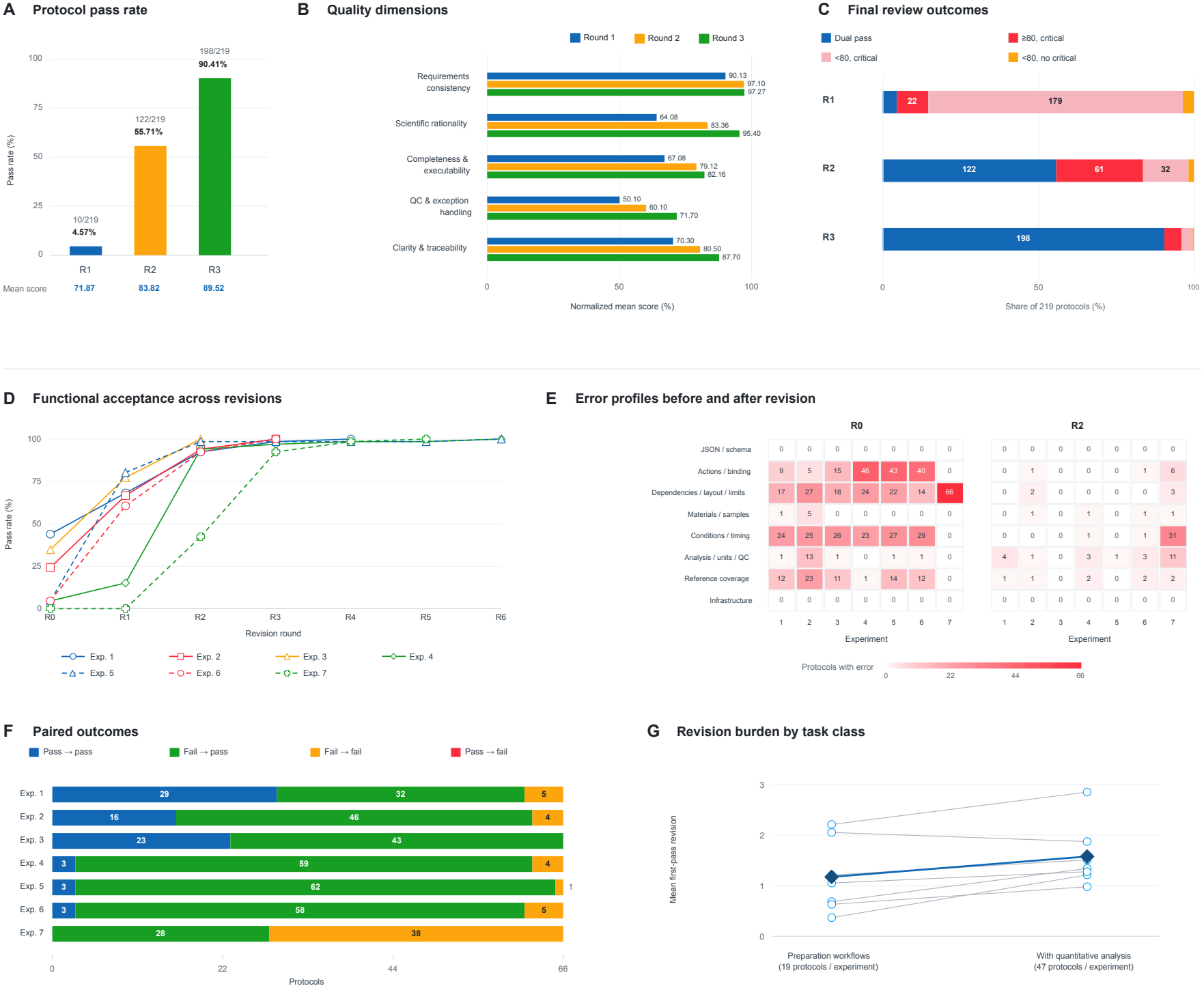
Dry-Lab Harness evaluation through protocol review and virtual validation. **(A)** Review pass rates over three rounds on the same 219 tasks; labels show passing counts, percentages, and mean composite scores. **(B)** Mean scores for five review dimensions, normalized to their maximum scores. **(C)** Review outcomes by round, separating protocols that passed both reviews from those with critical issues or scores below 80. Composite scores weight human assessment by two-thirds and *Review* agent assessment by one-third. **(D)** Virtual-validation pass rates over revisions across seven evaluations of the same 66 tasks. R0 denotes initial generation; only measured rounds are plotted. **(E)** Numbers of JSON-procedures with diagnostic issues by category at R0 and R2. A JSON-procedure can contribute to multiple categories, so counts are not additive. **(F)** Paired validation outcomes from R0 to R2 for each task within the same evaluation. **(G)** Mean revision counts to first pass for preparation workflows and tasks involving quantitative analysis, calculated separately for each evaluation. Open circles and connecting lines show the two task-class means for each of the seven evaluations; filled diamonds summarize their means. Panels D–G assess acceptance under the implemented simulation and software rules, not physical execution success.

Round 1 revealed a clear contrast: the normalized score for requirement consistency reached 90.13%, but the overall pass rate was only 4.57%. Scientific rationality scored 64.08%, completeness and executability scored 67.08%, and 201 of 219 protocols had at least one critical issue. Protocols could therefore follow the confirmed requirements and local resource constraints while still failing on scientific rationale, parameter settings, or step organization. Localized protocol generation is thus not only a matter of recognizing resource constraints; it requires integrating those constraints with experimental knowledge into a scientifically sound and operationally complete workflow.

In the next two rounds, the composite review pass rate rose to 55.71% and then 90.41%, while the mean score increased from 71.87 in round 1 to 83.82 and then 89.52. The number of protocols with critical issues fell from 201 to 93 and then to 20 (Fig. 2C). Representative critical issues across the three protocol-review rounds are summarized in Supplementary Table S8. In round 2, 61 protocols reached a score of 80 but still failed because of critical issues; in round 3 this number fell to 12. The increase in pass rate was therefore accompanied not only by higher scores but also by a marked reduction in issues that would prevent implementation.

Dimension-level changes showed which aspects improved most. Scientific rationality rose from 64.08% to 95.40%, the largest increase among the five dimensions, whereas requirement consistency reached 97.10% in round 2 and then remained stable. Later rounds therefore improved how the system made sound scientific decisions under the given conditions rather than merely increasing compliance with the input requirements. Completeness and executability, however, still reached only 82.16% in round 3, and quality control and exception handling remained the lowest dimension at 71.70%. Even when a protocol was scientifically sound, complete operating steps, quality-control requirements, and exception handling continued to limit review approval.

Taken together, the three rounds show that the Dry-Lab Harness can generate protocols that comply with local device, reagent, and consumable constraints, and that later rounds achieved higher scientific rationality, workflow completeness, and pass rates. These results support its ability to address the knowledge–environment mismatch at the planning level, but they do not by themselves demonstrate execution success on a real platform.

#### 3.1.2 Virtual-validation feedback improves procedure pass rates

We next evaluated feedback-guided procedure refinement seven times on the same 66 automation-eligible tasks. Each candidate JSON-procedure underwent simulated execution and software rule checking; failed candidates were revised using the returned status and errors. R0 denotes initial generation, and R1 and R2 denote the first and second revisions. These rounds measure repair within the evaluated tasks.

Averaged across the seven evaluations, the proportion of tasks that passed validation rose from 16.7% at R0 to 52.6% at R1 and 87.7% at R2 (Fig. 2D). The first round in which all 66 tasks passed varied from R2 to R6 across the evaluations (Supplementary Fig. S5). Under the given task set and validation conditions, virtual-validation feedback from the Wet-Lab Harness therefore allowed the model to repair procedures progressively and improve their compliance with local execution requirements.

Error analysis showed which constraints improved (Fig. 2E and Supplementary Fig. S5). Counting affected tasks separately in each evaluation, action- and binding-related diagnostics fell from 158 at R0 to 8 at R2; dependency, plate-position, and resource-limit diagnostics fell from 188 to 5; and condition- and timing-related diagnostics fell from 154 to 33. Revisions therefore mainly improved compliance with device-action invocation, object binding, and workflow dependencies. In contrast, analysis-, unit-, and quality-control-related diagnostics rose from 17 to 23, indicating that some downstream constraints still required attention in later rounds. Diagnostic counts were also affected by validation order: in the seventh evaluation, for example, all initial drafts were blocked at the resource-limit check, and timing and analysis diagnostics became visible only at R2. An increase in one diagnostic category may therefore reflect either the exposure of downstream problems after upstream ones were resolved or new problems introduced by revision, and must be interpreted together with the execution record.

Pairing R0 and R2 outcomes for each task within each evaluation showed that 328 of the 385 initial failures passed by R2 (85.2%). All 77 outcomes that passed at R0 remained passing at R2 (Fig. 2F). In the R0–R2 comparison, the increase in pass rate therefore came from repairing initially failing tasks, with no observed regression of initially passing tasks.

Improvement rates differed by task class (Fig. 2G). For each evaluation, we calculated the mean revision count to first pass within each task class. The median of these means was 1.05 rounds for preparation workflows and 1.34 rounds for tasks involving quantitative analysis. In six of the seven evaluations, tasks involving quantitative analysis took more rounds to pass on average. Quantitative analysis and quality-control tasks therefore generally required more rounds of feedback-driven revision than preparation workflows.

Within the defined execution environment, feedback from simulated execution and rule checking improved procedure pass rates and reduced several categories of execution-constraint problems. These results support feedback-guided protocol-to-procedure conversion as a basis for subsequent physical execution.

### 3.2 Physical execution feedback improves device orchestration and reusable operating policies

We evaluated two ways of using execution feedback to refine liquid handling (Fig. 3). On the Opentrons OT-2, we used measured runtimes to screen pipetting policies for plate replication and carried adopted policies into subsequent rounds. On the Tecan Freedom EVO 150, we used instrument logs to build a simulator for schedule search and evaluated scheduling revisions for both well rearrangement and sample normalization.

**Fig. 3.**
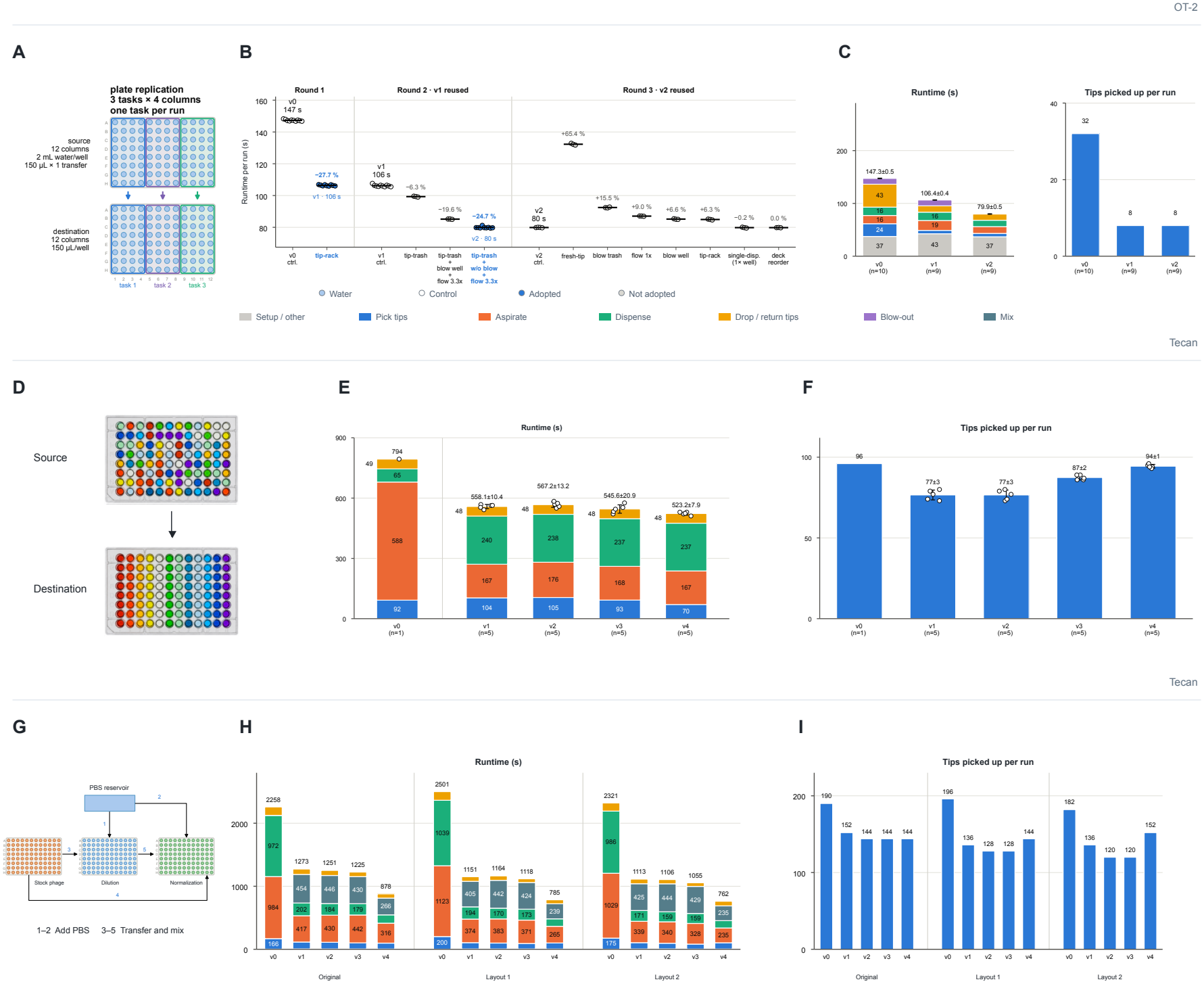
Feedback-guided refinement of liquid-handling policies and execution schedules. **(A–C)** OT-2 plate replication. (A) Source and destination well maps. Three tasks each span four columns and are executed in separate runs. (B) Runtime comparisons between controls and candidate policies across three rounds of refinement. Points represent individual runs, black horizontal lines indicate mean runtimes across the plotted runs, and blue denotes adopted policies. Percentages indicate the median paired change in runtime relative to the control for each round; negative values indicate shorter runtimes. Rounds 2 and 3 use the adopted v1 and v2 policies as their respective controls. Tip-rack and tip-trash denote tip reuse followed by return to the rack or disposal, respectively. (C) Runtime components and individual tips picked up per run for v0–v2. Runtime annotations report mean *±* standard deviation, with 10, 9, and 9 runs for v0, v1, and v2, respectively. **(D–F)** Well rearrangement. (D) Source deep-well plate before transfer (top) and destination microplate after transfer (bottom), with colored liquids illustrating the rearrangement of well contents. (E) Runtime and operation-level runtime components for v0–v4. (F) Individual tips picked up per run. Points represent layout-level values, and *n* denotes the number of layouts. Where shown, summary values report mean *±* standard deviation across layouts. **(G–I)** Phage titer normalization. (G) Workflow schematic. (H) Summed logged action times and their components for v0–v4 under the original layout (Original), Layout 1, and Layout 2. (I) Tip consumption per run. Each layout–version combination was executed once; no error bars are shown. Colors denote operation categories, as indicated in the legend.

#### 3.2.1 Pipetting policy refinement

On the liquid handler, the agent proposed changes to tip handling, aspiration grouping, flow rate, blowout, and transfer order for plate replication (Fig. 3A–C). The OT-2 plate-replication benchmark comprised three four-column tasks, with one task executed per run. In each round, the agent proposed policy changes and screened them against the current control in paired runs. For the two adopted policies, all paired comparisons exceeded the 10% improvement threshold. Comparisons used separate runs of the same task set.

The first adopted policy (v1) reused one set of tips across the four columns and returned it to the rack. Tip pickups fell from 32 to 8 per run, and the median paired runtime decreased by 27.7% relative to v0. In the next round, v1 served as the control. The adopted policy (v2) still used one tip set per run, discarded it after use, omitted blow-out, and increased the flow rate to 3.3 times the reference setting. Median paired runtime decreased by a further 24.7%, with tip pickups unchanged. In the third round, the best of seven variants was 0.2% faster than v2, below the 10% threshold, so v2 was retained. Across the two adopted rounds, the mean runtime fell from 147.3 s at v0 to 79.9 s at v2, a cumulative reduction of 45.8% (Fig. 3C).

#### 3.2.2 Well rearrangement scheduling

We used a 96-well rearrangement task to evaluate whether the Dry-Lab Harness could optimize liquid-handling schedules using physical execution feedback. As shown in Fig. 3D, an eight-channel LiHa on the Tecan Freedom EVO 150 liquid-handler transferred 12 types of colored liquids, randomly distributed across the source plate, to a destination plate according to a predefined well mapping, with a nominal transfer volume of 150 *µ*L per well. Throughout the experiment, the source-to-destination well mapping, plate layout, and hardware configuration were kept constant.

The Dry-Lab Harness includes a parameterized runtime simulator for the Tecan Freedom EVO 150 liquid-handler to predict the execution time of different scheduling strategies. The simulator was constructed based on instrument execution logs, from which action-level information was extracted, including tip pickup, aspiration, dispensing, mixing, tip disposal, and robotic arm movements. It parsed instructions in Gemini WorkList (GWL) format into ordered action sequences and estimated execution times based on action type, the number of active channels, arm position, and tip state. Using this simulator, the Dry-Lab Harness progressively explored and refined scheduling strategies, starting from the Tecan built-in strategy (v0) and generating four subsequent versions (v1–v4). The specific strategy changes in each version are summarized in Supplementary Table S1.

Runtime did not decrease monotonically across successive optimization versions (Fig. 3E). The mean runtime was 558.1 *±* 10.4 s for v1, increased to 567.2 *±* 13.2 s for v2, and subsequently decreased to 545.6 *±* 20.9 s and 523.2 *±* 7.9 s for v3 and v4, respectively. Compared with the single observed runtime of 794 s for v0, the mean runtime of v4 was 34.1% lower. Meanwhile, the mean number of individual tips picked up per run increased from 77 in v1 to 94 in v4 (Fig. 3F), showing that the shorter runtimes were not obtained by reducing tip consumption. These results show that iterative scheduling strategy refinement by the Dry-Lab Harness ultimately reduced the overall execution time of this task.

#### 3.2.3 Phage titer normalization scheduling

We further applied the Dry-Lab Harness to phage titer normalization to evaluate its scheduling optimization capability in a practical biological workflow. As illustrated in Fig. 3G, the workflow involved PBS addition, phage sample transfer, and subsequent mixing. To evaluate scheduling performance under different sample layouts, we used three layouts: Original layout, Layout 1, and Layout 2. The Original layout corresponded to the sample arrangement used in the subsequent phage titer normalization experiment, whereas Layout 1 and Layout 2 represented two alternative sample arrangements. Starting from the Tecan built-in strategy (v0), the Dry-Lab Harness progressively refined the scheduling strategy through four successive versions (v1–v4), with the specific modifications summarized in Supplementary Table S2. Each layout–version combination was executed once.

The effects of scheduling modifications depended on the sample layout (Fig. 3H). For example, following the introduction of cross-plate aspiration in v2, the summed logged action time decreased from 1273 to 1251 s under the Original layout and from 1113 to 1106 s under Layout 2, whereas it increased from 1151 to 1164 s under Layout 1. This indicates that the same scheduling modification can produce different execution outcomes under different sample layouts. Following subsequent strategy refinements, the summed logged action times in v4 decreased to 878, 785, and 762 s under the Original layout, Layout 1, and Layout 2, respectively, corresponding to reductions of 61.1%, 68.6%, and 67.2% relative to their respective v0 baselines. Meanwhile, tip consumption across the three layouts decreased by 24.2%, 26.5%, and 16.5%, respectively (Fig. 3I).

Overall, the Dry-Lab Harness improved execution efficiency across all three sample layouts through successive scheduling strategy refinement. These results demonstrate both the influence of sample layout on scheduling performance and the capability of the Dry-Lab Harness to optimize liquid-handling schedules under different layout conditions, providing efficient automated execution support for the subsequent phage ELISA experiment.

### 3.3 Experimental outcomes guide method revision and biological screening

To examine the protocol–outcome mismatch, we trace how biological observations informed vector and assay development, then report the resulting workflow’s batch performance. Of 96 starting construction samples, 93 yielded a sequence-correct clone. These clones were carried forward to titer measurement with two additional controls, titer-guided normalization, and ELISA screening. Workflow run completion and sample-level biological success are reported separately.

#### 3.3.1 Molecular cloning

To establish a negative-selection vector for high-throughput automated molecular cloning, the SEA-Lab system analyzed the relevant literature. Drawing on the use of a replaceable *ccdB* cassette for background suppression by Qian et al. [14], it proposed a *ccdB* module for the existing phage-display vector pDWPD, yielding the pDWPD V1– V8 series, whose promoter–RBS combinations and promoter sequences are shown in Fig. 4A and Supplementary Fig. S7A. Construction of the variants was attempted by an external supplier in the *ccdB* -tolerant host *E. coli* DB3.1; plasmids recovered successfully were then transformed into *E. coli* TG1 to evaluate negative selection (Fig. 4B and Supplementary Fig. S7B). To control for batch-to-batch variation in transformation efficiency, TG1 transformations of pDWPD V1, pDWPD V2, and pDWPD V8 each included the corresponding original plasmid as a matched positive control.

**Fig. 4.**
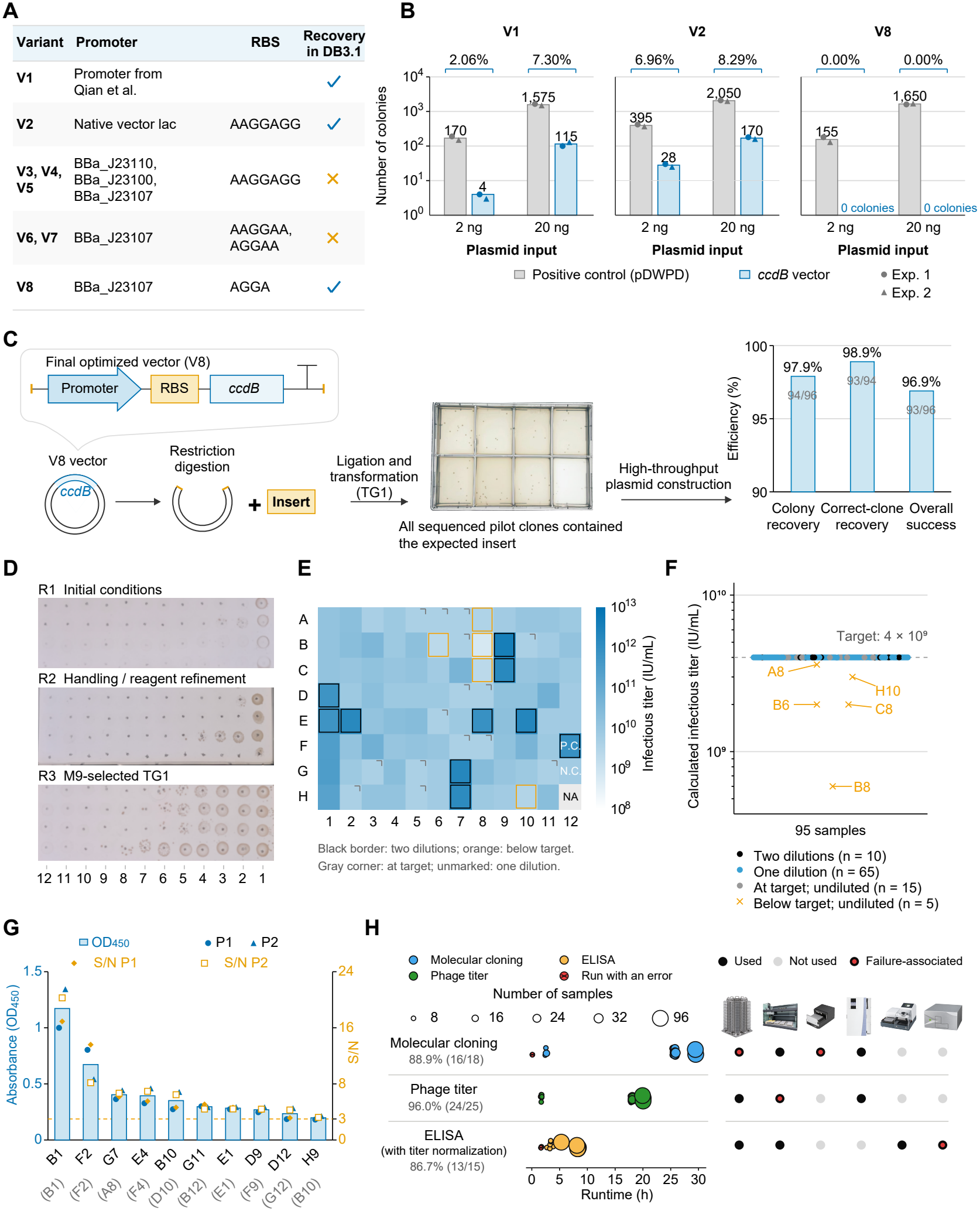
Phage-display construction, titer normalization, ELISA screening, and automated workflow performance. **(A)** Promoter–RBS combinations and vector recovery for pDWPD V1–V8 in *E. coli* DB3.1. Check marks indicate recovery of the expected vector. **(B)** TG1 colony counts after transformation with 2 or 20 ng of the indicated vectors. Bars show means of two experiments, symbols show individual counts, and percentages are relative to the matched positive control using unrounded means. **(C)** Cloning with pDWPD V8, representative plates from an eight-construct pilot, and outcomes of the subsequent 96-construct batch. **(D)** Three rounds of phage infection titer-assay development. Dilutions run from 10^*−*12^ to 10^*−*1^, left to right. **(E)** Infection titers by well on a log scale (IU/mL). Black outlines indicate two-step dilution, gray corners indicate samples at target, and orange outlines indicate samples below target. P.C. and N.C. denote positive and negative controls. **(F)** Calculated pre-blocking titers for 95 samples, including controls. The dashed line marks the target of 4 *×* 10^9^ IU/mL, and orange labels identify the five below-target samples. **(G)** Ten clones with *S/N >* 3 on both plates, ranked by mean OD_450_. Blue bars and symbols show mean and plate-specific OD_450_, respectively. Orange symbols show plate-specific *S/N* on the right axis. Parentheses give original plate positions. **(H)** Runtime and device use for cloning, titer measurement, and ELISA with normalization. Each bubble is one run, sized by sample count. From left to right, the device icons represent the labware carousel, liquid handler, thermal cycler, incubator, microplate washer, and microplate reader. Gray text reports completion rates. Dark and light gray dots indicate devices used and unused, respectively. Red centers mark runs with errors or devices associated with failures.

Initial tests showed that pDWPD V1, which uses the initial *ccdB* module, and pDWPD V2, which combines the existing *lac* promoter with the RBS sequence AAGGAGG, could both be constructed in DB3.1, but both still produced background colonies after transformation into TG1, indicating that negative selection was insufficient to suppress background clones completely (Fig. 4A,B and Supplementary Fig. S7B). The SEA-Lab system then combined the constitutive Anderson promoters BBa J23110, BBa J23100, and BBa J23107 [15] with the RBS sequence AAGGAGG to generate pDWPD V3– pDWPD V5. Despite repeated construction attempts, none of these variants reproducibly yielded the intended plasmid in DB3.1. Some recovered clones contained base deletions or other sequence variants in the promoter or regions adjacent to *ccdB* (Supplementary Fig. S7C). These observations suggest that the promoter–RBS combination may influence recovery of the intended constructs.

Subsequent optimization therefore considered two conditions simultaneously: successful construction in DB3.1 and suppression of background colony formation in TG1. Keeping the BBa J23107 promoter, the SEA-Lab system progressively adjusted the RBS sequence. pDWPD V6 and pDWPD V7, using AAG-GAA and AGGAA, respectively, still could not be recovered as intact plasmids, whereas pDWPD V8, using AGGA, was successfully constructed in DB3.1 (Fig. 4A). Functional validation showed that transformation of TG1 with 2 ng or 20 ng of pDWPD V8 produced no colonies, whereas the matched positive controls formed clear colonies (Fig. 4B and Supplementary Fig. S7B). Under these conditions, the BBa J23107–AGGA–*ccdB* architecture therefore combined recovery of intact plasmids in DB3.1 with no detectable colony formation in TG1.

To evaluate cloning with pDWPD V8, the *ccdB* cassette was excised by NheI and NotI digestion and replaced with a primer-annealed insert, followed by transformation into chemically competent TG1 cells (Fig. 4C). In an independent pilot experiment involving eight constructs, all eight yielded colonies, and all picked clones submitted for sequencing contained the expected insert (Fig. 4C). These results supported subsequent evaluation in a separate 96-construct batch.

The cloning workflow was integrated into the SEA-Lab automation platform. The system coordinated the robotic arm, liquid handler, thermal cycler, incubator, and colony picker to perform oligonucleotide annealing, ligation, transformation, colony picking, and cultivation (Supplementary Fig. S6A). Of 18 automated runs, 16 completed the full workflow, corresponding to a completion rate of 88.9% (16/18; Fig. 4H). The two failures involved incorrect positioning at the thermal cycler and an offset in the transfer position of the labware carousel (Fig. 4H). A complete 96-well batch required approximately 30 h from workflow initiation to completion (Fig. 4H and Supplementary Fig. S6D). These results demonstrate the feasibility of integrating the complete workflow and processing samples in parallel on the platform.

In the 96-sample construction batch, colonies were recovered for 94 samples, giving a colony recovery rate of 97.9% (94/96); only A7 and H10 failed to produce colonies. Two clones were selected for sequencing from each colony-positive sample, except F3, for which only one clone was available. At least one sequence-correct clone was recovered for 93 samples, whereas the sole F3 clone did not match the expected sequence. The correct-clone recovery rate among colony-positive samples was therefore 98.9% (93/94), and the overall construction success rate across all starting samples was 96.9% (93/96). One sequence-correct clone from each successful construct was retained and transferred to the final phage culture plate (Fig. 4C and Supplementary Fig. S7D,E).

Together, vector screening, clone validation, and parallel construction showed that pDWPD V8 could be recovered intact from the construction host while effectively suppressing background colonies in TG1, and that replacement of *ccdB* by the target fragment supported recovery of correct clones. In this batch, 93 sequence-correct clones were obtained for the subsequent phage-display workflow.

#### 3.3.2 Phage infection titer measurement

To establish an infection-titer assay suitable for high-throughput phage screening, we infected TG1 with phage, counted resistant colonies, and optimized sample handling and host-cell state over three rounds (Fig. 4D). Under the initial conditions, most spot positions showed no visible colony growth, making it difficult to obtain counts for titer calculation. Adjustment of dilution and mixing and replacement of the relevant reagents improved colony growth for some samples, but most samples still lacked a clear dilution gradient, indicating that these operational changes were insufficient. Because loss of the F factor in TG1 could affect phage infection, TG1 selected on M9 medium was used in the next round. In the third round, colony growth and its distribution across dilutions were clearer, and dispersed colonies were visible in some spot regions, providing a basis for choosing an appropriate dilution and calculating infection titer. This improvement suggested that host-cell state was an important determinant of assay performance, although it did not directly establish that the earlier anomalies were caused by F-factor loss.

Phage preparations derived from the 93 sequence-correct clones were subsequently analyzed using the optimized infection-titer assay. To connect with subsequent high-throughput operations, samples were re-arrayed, with positive and negative controls in F12 and G12, respectively, and H12 left blank (Supplementary Fig. S7E). Infection titers of different samples spanned several orders of magnitude, indicating substantial variation in the concentration of infectious phage after packaging and providing a quantitative basis for adjusting sample input in the subsequent phage ELISA (Fig. 4E).

To assess the method in multi-device execution, we evaluated the automated phage-titer workflow, which the labware carousel, liquid handler, and incubator completed jointly, linking sample dilution, spot plating, and cultivation (Supplementary Fig. S6B,E). A complete 96-well batch required approximately 20 h from workflow initiation to completion. Of 25 recorded runs, 24 completed, giving a workflow completion rate of 96.0% (Fig. 4H). The main recorded failure involved an inappropriate spot-plating height on the liquid handler, which caused the plate to be lifted during operation, showing how the geometry between the pipetting action and the plate position affects stable workflow execution. These results span assay optimization, titer quantification, and automated workflow execution, showing how the infection-titer assay was connected to a multi-device pipeline.

#### 3.3.3 Phage ELISA

Before ELISA, 95 phage preparations, comprising 93 sequence-correct constructs and two controls, under-went titer-guided input adjustment toward a pre-blocking target of 4 *×*10^9^ IU/mL. Of these, 65 were diluted directly, and 10 required an intermediate dilution to maintain a minimum phage-transfer volume of 2 *µ*L. Fifteen preparations already at the target and five below it were used undiluted (Supplementary Fig. S7G). Fig. 4F shows final titers calculated from the measured stock titers and the pipetting volumes specified in the worklists, rather than titers independently measured after dilution. The five below-target preparations therefore retained lower nominal phage inputs. All preparations were pre-blocked before the binding step. Well identifiers in the titer and ELISA analyses refer to the re-arrayed phage culture plate (Supplementary Fig. S7E).

The 93 clones were screened in duplicate across two plates, each containing positive, negative, and blank controls. Readings were matched by well, and the OD_450_ distributions for both plates and their means were displayed using a common color scale (Supplementary Fig. S7H). The Pearson correlation between plates was 0.928, and the Spearman rank correlation was 0.643, indicating a positive association between measurements.

*S/N* was calculated separately for each plate as the sample OD_450_ divided by the negative-control OD_450_ in G12 on the same plate, without blank subtraction. Ten clones—B1, F2, G7, E4, B10, G11, E1, D9, D12, and H9—exhibited *S/N* values greater than 3 on both plates and were ranked by mean OD_450_ for display (Fig. 4G). Signals from the five below-target preparations should also be interpreted in light of their lower phage inputs, rather than attributed directly to weaker binding.

This primary screen established a prioritized set of ten candidate clones, with B1, F2, G7, and E4 consistently ranking highest across both plates, providing a basis for subsequent characterization of binding specificity and signal stability (Fig. 4G).

Under SEA-Lab system control, the automated ELISA workflow coordinated the labware carousel, liquid handler, microplate washer, and microplate reader. Sample-specific worklists guided titer normalization, while the assay workflow comprised coating, blocking, phage binding, detection-antibody incubation, substrate addition, and reaction termination, with intervening washes. Plate transfers and device operations followed task dependencies, concluding with absorbance measurement at 450 nm. Of 15 recorded runs, 13 completed the full workflow (86.7%). ELISA run failures involved microplate reader disconnection and stalling (Fig. 4H). A complete 96-well batch, including titer normalization, required approximately 8 h. Detailed workflow steps, operating parameters, and a representative execution timeline are provided in Supplementary Fig. S6C,F.

## 4 Discussion

SEA-Lab establishes an experiment-grounded feedback architecture in which local constraints shape experimental planning, physical execution informs operational policies, and measured outcomes guide experimental strategy. The three mismatches — knowledge–environment, plan–execution, and protocol– outcome — define distinct requirements for autonomous experimentation: locally feasible plans, verified execution, and scientifically interpretable outcomes. Across protocol review, virtual validation, instrument execution, and biological application, the results demonstrate how multi-level feedback translates AI-generated plans into executable experiments and enables laboratory experience to inform subsequent refinement.

Addressing the knowledge–environment mismatch required integrating resource awareness with scientific reasoning and operational detail. High initial requirement-consistency scores paired with a low overall pass rate showed that recognizing local devices and materials was insufficient to generate acceptable experimental plans. Local constraints had to be incorporated into scientifically sound methods, complete operating steps, and appropriate quality controls. Iterative protocol review supported this integration at the design level, whereas virtual validation translated errors in device interfaces, task dependencies, and resource configurations into structured diagnostics for targeted procedure repair before physical runs. These complementary feedback mechanisms connected scientific planning with the explicit requirements of the local execution environment.

Addressing the plan–execution mismatch required physical runtime observations, which revealed how execution strategies depend on instrument dynamics and plate layouts. Greater tip reuse did not consistently shorten runtime in well rearrangement, and identical scheduling changes produced opposite effects across initial layouts in phage titer normalization. Evaluating operational changes within complete work-flows was therefore essential. Combining action-level instrument logs with simulation linked observed device behavior to revisions of timing estimates and candidate schedules. Adopted pipetting policies then became the reference for subsequent optimization rounds, demonstrating that an improvement could be retained and used as the starting point for further refinement. Execution feedback thus supported both adaptation to the physical platform and continued optimization from previously established policies.

The biological application extended this feedback process to the protocol–outcome mismatch, showing why completed device execution must be evaluated alongside biochemical and functional readouts. Experimental observations redirected scientific decisions across distinct levels: in negative-selection vector engineering, the failure to recover intact plasmids for intermediate promoter–RBS designs required balancing construct recoverability in DB3.1 against background suppression in TG1; in phage titration, persistent gradient anomalies despite liquid-handling adjustments shifted attention from operational parameters to host-cell preparation. These cases illustrate how experimental outcomes can identify a need for method-level revision beyond changes to the execution procedure. Integrating the resulting methods across molecular cloning, phage titration, titer-guided normalization, and ELISA screening demonstrated multi-stage coordination while separating workflow run completion from sample-level biological success.

The three mismatches locate improvement at different levels of the experimental process: the integration of local constraints with scientific reasoning, the calibration of execution models and policies, and the revision of methods in response to measured outcomes. SEA-Lab connects these levels by making execution policies and experimental strategies explicit objects of feedback-guided revision and reuse. Within-task repair corrects the current procedure, while retained policies and strategies allow experience to inform later planning. The adopted-policy comparisons demonstrate this reuse across optimization rounds, and the biological cases show how experimental outcomes can redirect method-level decisions. Together, these capabilities establish a concrete architecture for experiment-grounded recursive self-improvement.

## 5 Limitations

This study was conducted within a defined laboratory environment, device configuration, and task set, providing a basis for examining the relationships among local resource constraints, protocol generation, and physical execution. Iterative protocol-generation evaluations used the same task set, and automation tasks were selected for compatibility with local device capabilities. The current results therefore reflect SEA-Lab’s performance under the evaluated conditions. Evaluations on independent task sets and deployment across alternative platforms would further assess its adaptability to new tasks, device configurations, and laboratory environments. Such studies could also identify which knowledge and execution policies transfer across environments and which require local reconfiguration, informing broader deployment.

The current evaluation demonstrated feedback-guided protocol revision and policy optimization and provided an initial assessment of adopted-policy reuse in subsequent rounds. These results offer a practical basis for investigating how laboratory experience can inform later planning. Further work should examine whether and how improvements achieved within individual optimization cycles translate into sustained performance gains as experience accumulates. Longer execution sequences and targeted controlled comparisons would help distinguish the contributions of policy reuse, repeated feedback, and search, while assessing the continued applicability of retained experience across successive tasks. Future evaluations should jointly consider efficiency, operational quality, and execution stability to characterize the effects of policy improvement and guide the development of longer-term experience accumulation and reuse.

The biological application demonstrated the integration of multiple experimental stages and their feasibility in the evaluated sample batch, illustrating how planning, device execution, and outcome feedback can be connected in practice. Broader experimental applicability and the primary-screen results require further confirmation through independent replication and subsequent characterization. Human participation remains necessary for experimental preparation, protocol review, and certain externally performed experimental steps, while autonomous operation is bounded by available device interfaces and resources. Broader device coverage, more comprehensive experimental records, and automated quality-control feedback could extend the range of tasks performed autonomously. These developments would support further evaluation of continuous operation, autonomous coordination, and outcome-guided decision-making in more complex experimental workflows.

## Supplementary information

The Supplementary Information includes Figs. S1–S7, Tables S1–S8, and Sections S1–S5.

## Author contributions

C.W. and Y.J. conceived and designed the study. R.G., B.Z., Y.J. and H.C. implemented the code, performed the evaluations, and analyzed the data. B.Z. and T.Z. integrated and tested the instruments. X.W., Y.G., M.F., X.M., Z.H. and Y.H. participated in the automated biological experiments. Y.J., R.G., H.C., X.W. and Y.G. drafted the manuscript. Q.W. reviewed and edited the manuscript. Y.G. and C.W. supervised the study. All authors read and approved the final manuscript.

