## Supplementary Information for "SEA-Lab: Experiment-grounded recursive self-improvement in an automated laboratory"

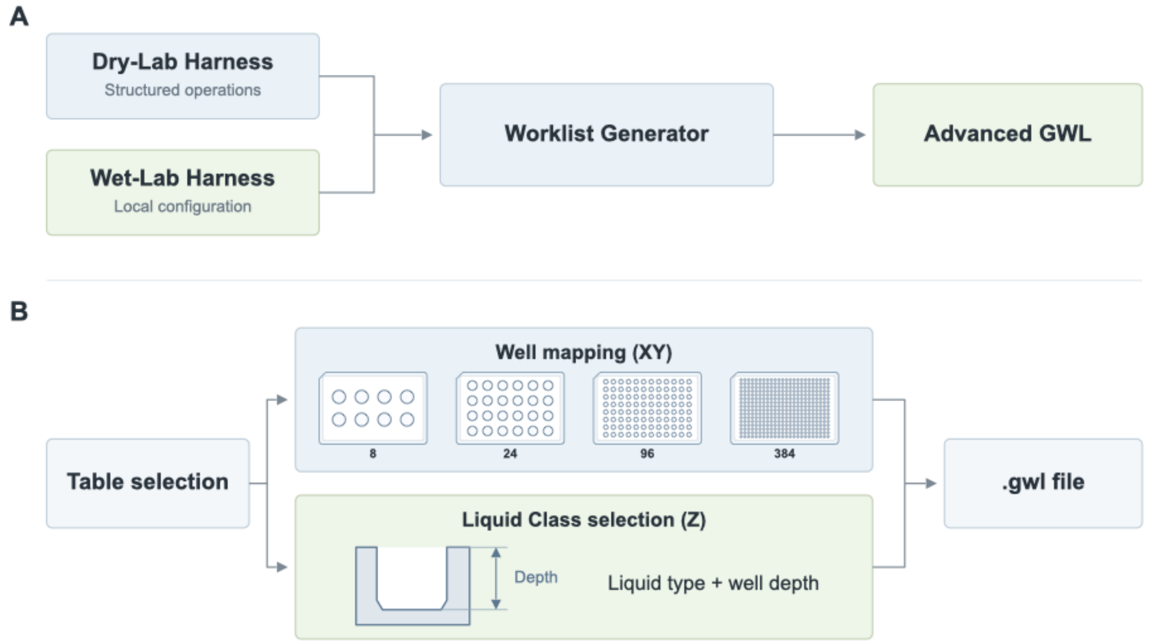


**Supplementary Figure S1. Worklist generation using local laboratory configurations.** The Worklist Generator combines structured experimental operations with local execution configurations to generate advanced worklists for the Tecan Freedom EVO 150. It selects a predefined deck layout and maps operation parameters to registered labware and EVOware Liquid Class entries. XY addressing uses geometry classes for 8-, 24-, 96-, and 384-well labware, together with predefined access points and 15 deck layouts. Z-related liquid handling uses registered Liquid Class entries selected by liquid type and well-depth class. These entries provide the aspiration, dispensing, and mixing settings used by the workstation.

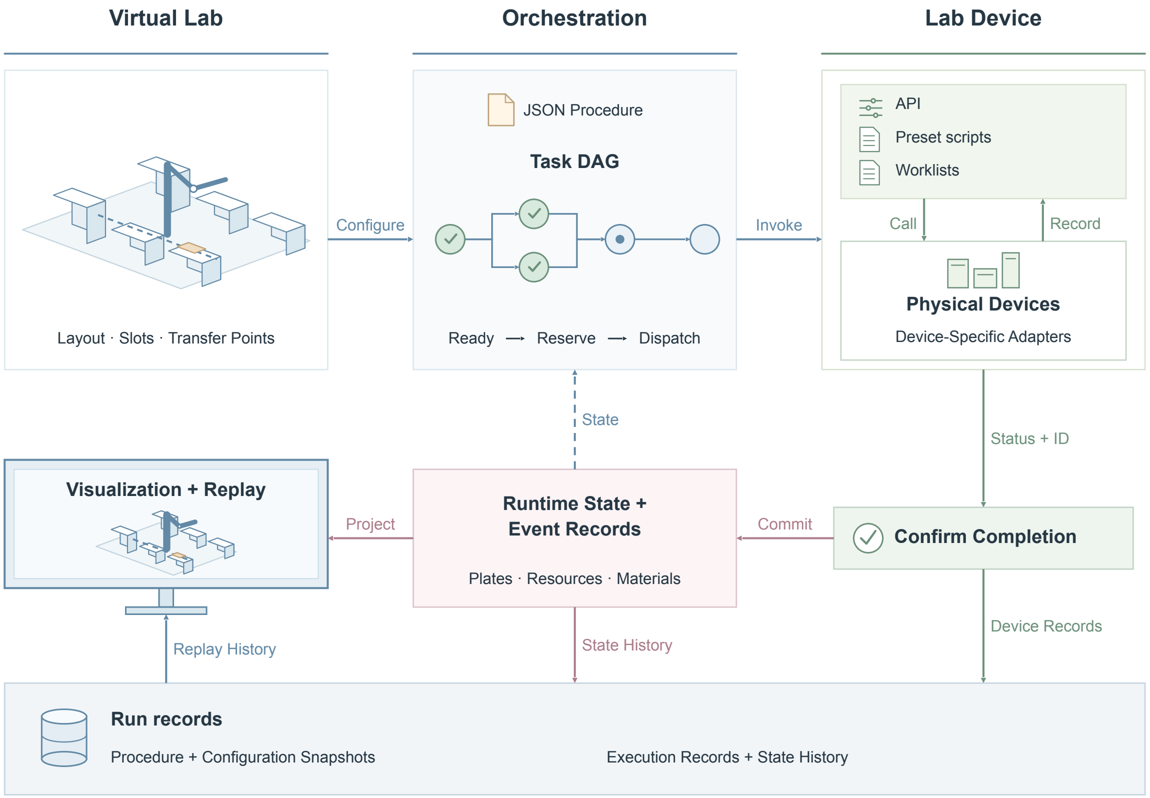


**Supplementary Figure S2. Wet-lab harness architecture for device execution and run-record storage.** The orchestration layer parses each JSON-procedure into a directed acyclic task graph. Virtual Lab provides device layouts, logical slots, and transfer points. Lab Device dispatches eligible tasks through device-specific adapters using parameterized actions, named programs, or worklists. Execution identifiers and device records support completion verification and runtime-state updates. Persistent storage retains procedure and configuration snapshots, execution records, and labware, resource, and material histories. Shared task, device, plate, slot, and execution identifiers link spatial configuration, device calls, and execution replay.

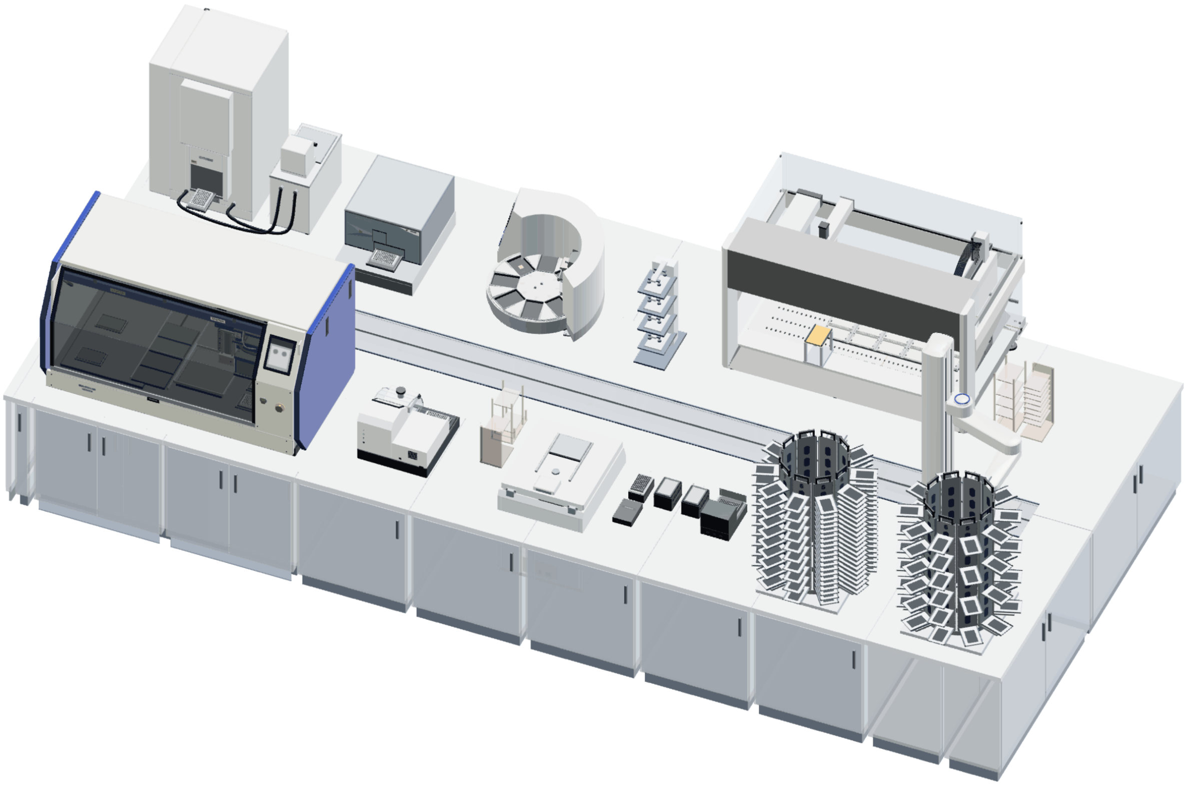

**Supplementary Figure S3. Physical layout of the automated laboratory platform.** Three-dimensional rendering of the instruments, labware storage units, and robotic plate-transfer system used for device coordination and labware movement.

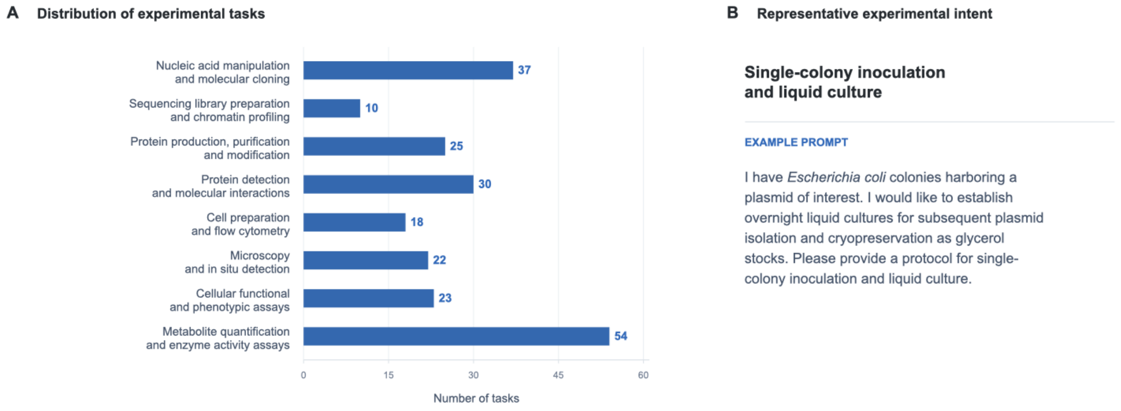


**Supplementary Figure S4. Task coverage and a representative input for the protocol-generation benchmark.** (A) Distribution of the 219 tasks across eight experimental domains. Each task was assigned to one primary domain; labels indicate task counts. (B) Representative experimental intent for single-colony inoculation and liquid culture, showing how an experimental objective and requested output are supplied to protocol generation.

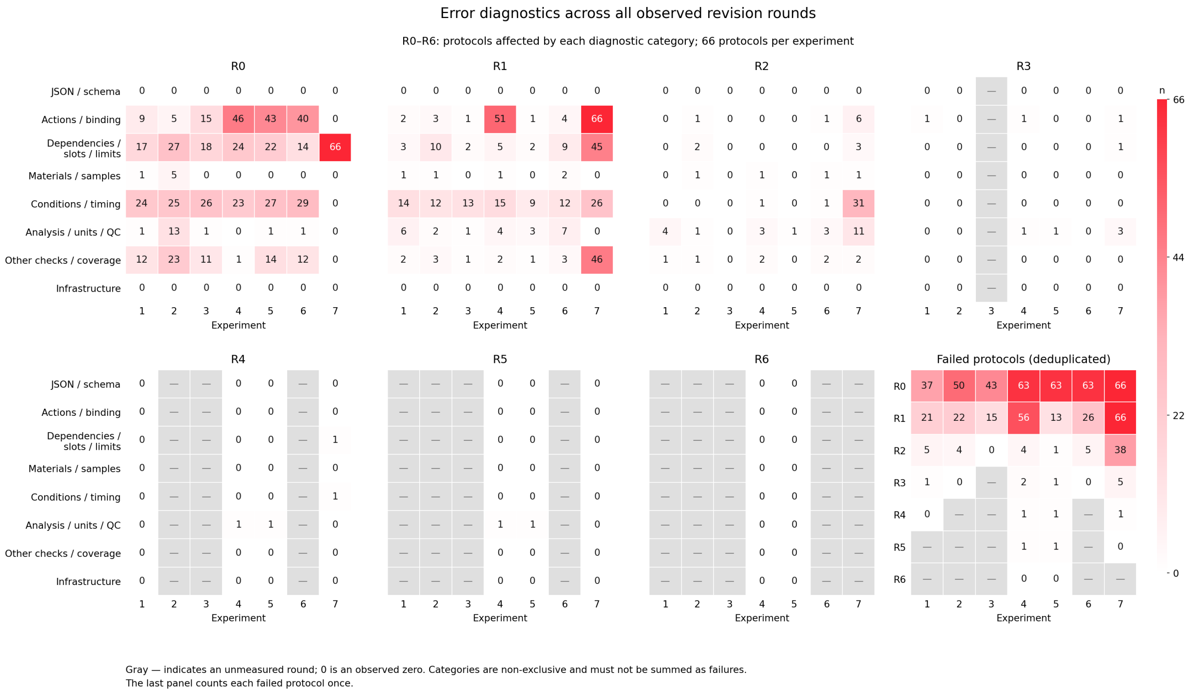


**Supplementary Figure S5. Diagnostic categories across iterative virtual validation.** Experiments 1–7 are repeated evaluations of the same 66 automation-eligible tasks. R0 denotes initial JSON-procedure generation, and R1–R6 denote successive revisions. Each category heatmap reports the number of procedures with at least one diagnostic in that category. A procedure is counted once per category and can appear in several categories. The lower-right heatmap reports failed-procedure counts deduplicated across categories. Cell values and red intensity indicate counts. Gray cells marked with a dash denote unmeasured rounds; zero denotes an observed count of zero.

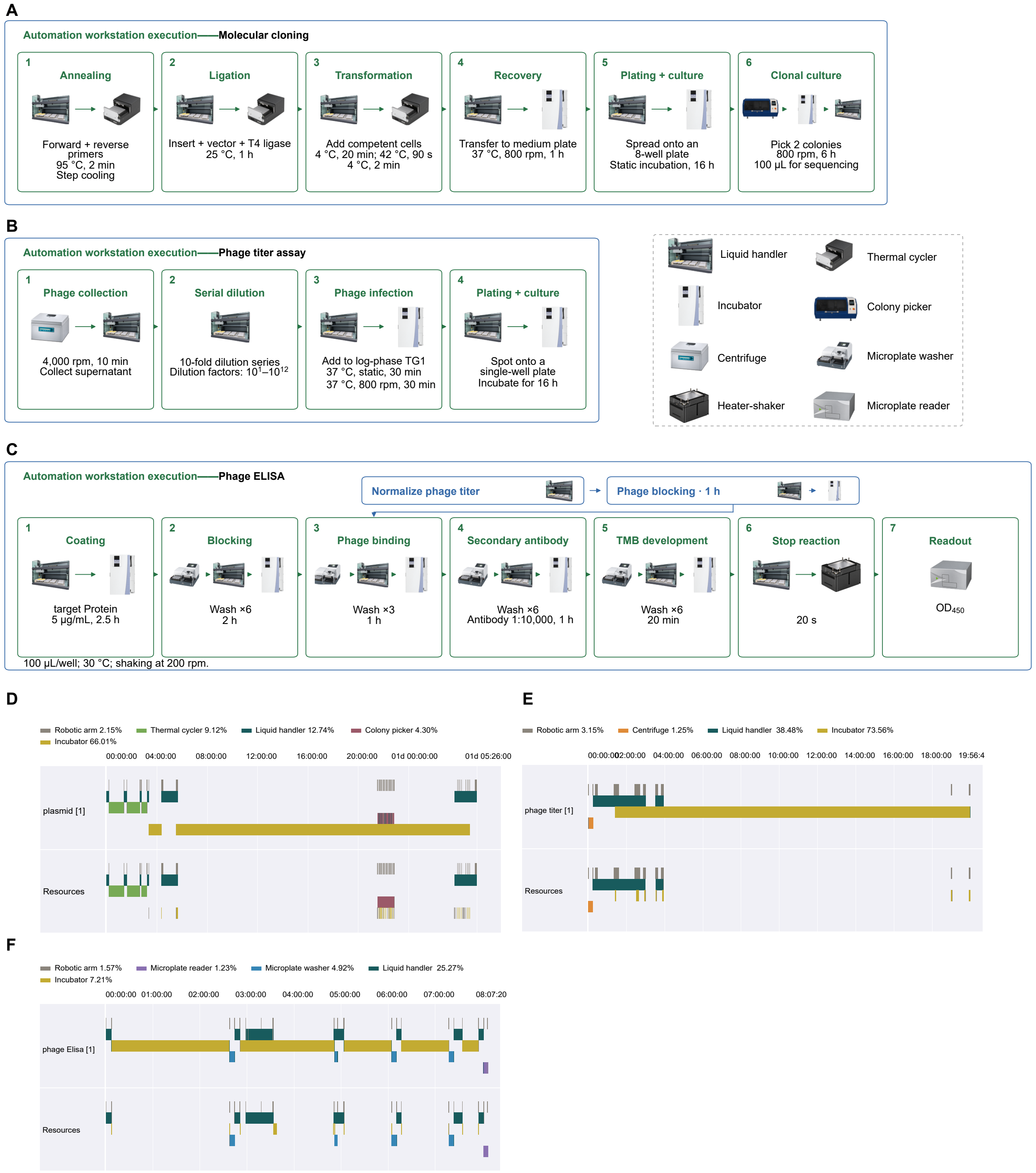


**Supplementary Figure S6. Automated workflows and execution timelines. (A–C)** Workflows for molecular cloning, phage titer measurement, and phage ELISA, respectively. Rounded boxes denote stages, and arrows indicate transfers between devices. Labeled times indicate incubation in the thermal cycler or incubator, or the duration of centrifugation. **(D–F)** Representative execution timelines for the same workflows, with approximate durations of 30 h, 20 h, and 8 h, respectively. Device colors are consistent across workflows. Percentages in the legend show device utilization reported by the software.

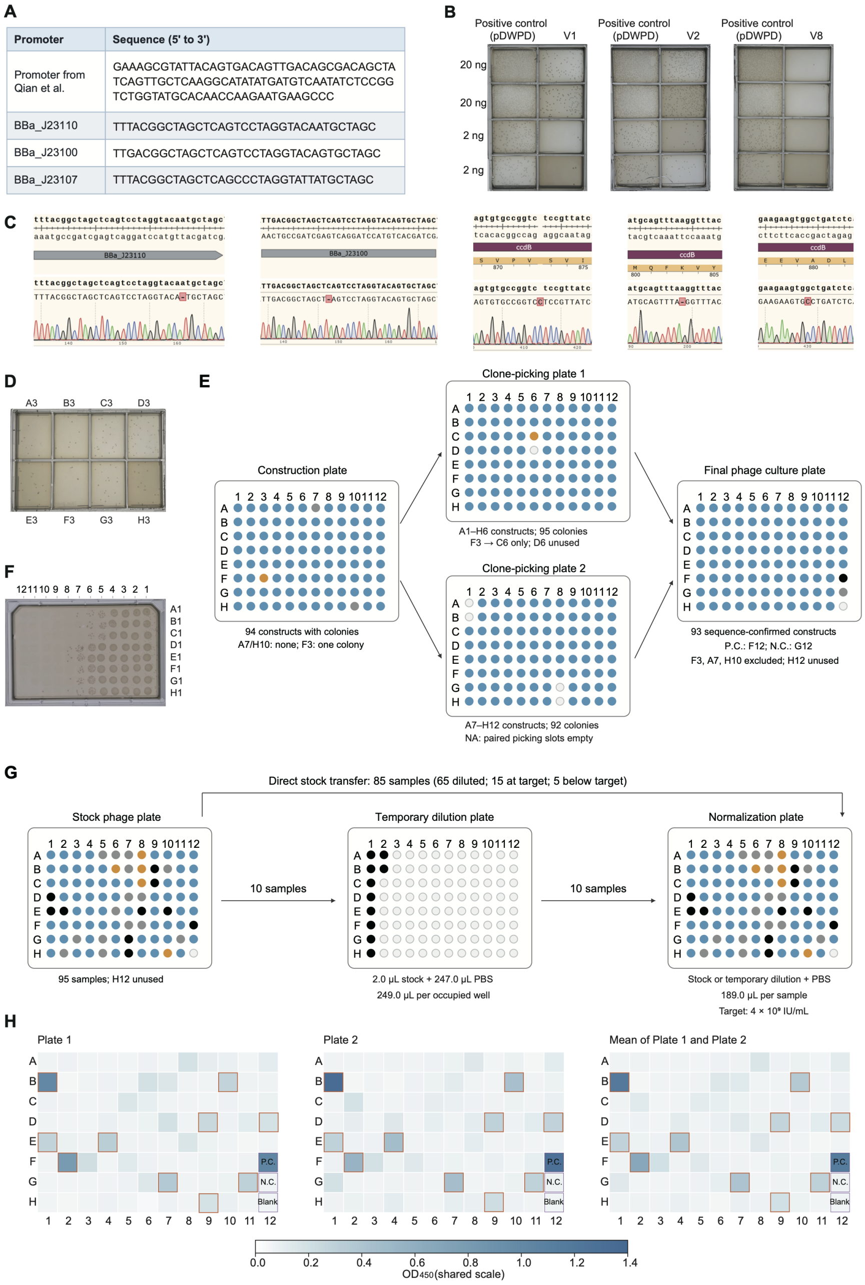


**Supplementary Figure S7. Vector development, sample re-arraying, and phage ELISA screening. (A)** Promoter sequences used in vector development, shown in the 5′ to 3′ direction. **(B)** Representative plates after transformation of TG1 with pDWPD_V1, pDWPD_V2, or pDWPD_V8 and the matched original vector as a positive control. Two experiments are shown for each condition at 2 and 20 ng plasmid inputs. **(C)** Representative sequence variants observed during construction of pDWPD_V3–pDWPD_V7. Reference sequences, alignments, and Sanger traces show the promoter or *ccdB* region. Red marks indicate sequence differences. **(D)** Representative plates for A3–H3, the eight samples in column 3 of the 96-sample construction batch. **(E)** Clone picking and re-arraying. A7 and H10 yielded no colonies. Up to two colonies were picked per colony-positive construct, with only one available for F3. Blue circles indicate colonies or retained samples, orange circles identify F3 and its clone, and gray or open circles indicate unavailable or unused positions. The re-arrayed plate contained 93 sequence-confirmed clones, with the positive and negative controls in F12 and G12, respectively. H12 was unused. **(F)** Representative spot plates from serial tenfold dilutions. Sample well identifiers are shown at right and dilution labels above the plate images. **(G)** Titer-guided normalization before blocking. At the target of $4 \times 10^9$ IU/mL, 65 preparations were diluted directly (blue), 10 used an intermediate dilution (black), 15 were already at the target (gray), and five below the target were used undiluted (orange). Intermediate dilutions used 2 $\mu$L stock and 247 $\mu$L PBS per well. The final volume per preparation was 189 $\mu$L. Open circles indicate unused wells. P.C., N.C., and Blank denote positive, negative, and blank controls. **(H)** Raw $OD_{450}$ for two ELISA plates and their mean, displayed with a common color scale. Orange outlines identify B1, F2, G7, E4, B10, G11, E1, D9, D12, and H9. P.C., N.C., and Blank denote positive, negative, and blank controls.

### S1 Dry-Lab Harness for worklist development

#### S1.1 Worklist generation and local configuration

The Worklist Generator translates structured experimental operations into Advanced GWL worklists for the Tecan Freedom EVO 150 (Supplementary Fig. S1). Inputs specify source and destination wells, liquid type, transfer volume, and mixing requirements. The wet-lab harness supplies predefined deck layouts, labware and position bindings, inventory state, and registered EVOware Liquid Class entries. The generator selects a compatible layout and compiles instructions that preserve the specified task and source–destination mapping. Deck layouts are represented as Tables in the local EVOware configuration.

Well addressing uses registered geometry classes for 8-, 24-, 96-, and 384-well labware. Fifteen predefined layouts cover the nonempty combinations of these four classes. Reservoirs with non-standard geometry use predefined access-point mappings to compatible well templates. These mappings provide a common addressing framework for the configured labware.

Liquid-handling settings are supplied by EVOware Liquid Class entries selected according to liquid type and the well-depth class D1–D4 at the operation location. Aspiration, dispensing, and mixing use entries associated with the relevant source, destination, or mixing location. Physical settings, including tip Z positioning, remain in the registered device configuration. Worklist compilation applies the operating rules of the relevant arm and the geometry mappings configured for each labware type.

#### S1.2 Runtime simulation and well rearrangement scheduling

Schedule search for the 96-well rearrangement task varied source-well visit order, channel assignment, destination dispense order, and tip management while preserving the specified well mapping and transfer volumes. Candidate schedules were compiled into GWL worklists and checked for consistency in well mapping, transferred volume, and liquid held in each channel.

A parameterized simulator decoded each worklist into an ordered action sequence and estimated runtime from action type, active channels, arm position, and tip state. Time-stamped instrument logs supplied action-level timing for calibration and feedback for subsequent revisions. Supplementary Table S1 summarizes the well-rearrangement versions; Supplementary Table S2 summarizes the cumulative changes used in titer-normalization scheduling.

**Supplementary Table S1.** Schedule revisions for the Tecan well rearrangement task.

| Version | Scheduling change | Interpretation |
|---|---|---|
| v0 | Tecan's native protocol. | Built-in Tecan cherry-picking strategy. |
| v1 | Searched source-column order, channel assignment, and execution scheduling using the cost of the compiled GWL. | Optimization evaluated executable instruction sequences. |
| v2 | Added candidates with liquid retained across transfer groups and asynchronous re-aspiration. | Expanded the search space; measured runtime did not improve over v1. |
| v3 | Jointly searched source-column order and channel assignment. | Multiple scheduling decisions were adjusted together. |
| v4 | Adjusted retention of empty tips or replacement with fresh tips and added preparation of fresh tip boxes. | Liquid-handling instructions followed v3 while the resource strategy changed. |

**Supplementary Table S2.** Schedule revisions for the phage titer normalization task.

| Version | Key Modifications | Rationale |
|---|---|---|
| v0 | Tecan's native protocol. | Tecan's native strategy. |
| v1 | Adopted a multi-dispense strategy: PBS was aspirated once using the 8-channel pipetting head and then dispensed across multiple wells. | Reduced repeated PBS aspiration. |
| v2 | Introduced a cross-plate aspiration strategy, with aspiration performed separately from the Phage and Dilute plates. | Improved channel utilization. |
| v3 | Returned used tips to their original rack positions. Once all tips had been used, the entire rack was moved to the disposal position. | Shortened the travel path to the remote tip-disposal area. |
| v4 | Combined transfers within the same column and performed mixing locally in the destination wells immediately after dispensing. | Reduced within-column splitting and repeated visits to destination wells. |

#### S2 Protocol to JSON conversion prompt interface

Protocol-to-JSON conversion combines the source protocol and experimental objective with confirmed user facts, registered device capabilities, initial inventory, and the platform schema. Supplementary Note summarizes the interface used to generate, validate, and revise a procedure. Experiment-specific prompts are recorded for the corresponding experiments.

**Supplementary Note.** Prompt template for protocol-to-JSON conversion.

```
Task:
Generate a JSON-procedure for the given protocol and wet-lab harness environment that
    conforms to the platform schema.

Inputs:
1. Source protocol and experimental objectives.
2. Confirmed user-provided facts and experimental conditions.
3. Registered devices, actions, and parameter constraints.
4. Labware definitions, initial positions, and inventory.
5. JSON Schema and the tools allowed for this run.

Generation:
Use registered actions and declare required dependencies.
Preserve the experimental objectives, operation order, and confirmed conditions.
Report missing information and unsupported operations explicitly.
Do not invent device capabilities, inventory, parameters, or completion evidence, or
    use a wait task or an unverified external completion as a substitute.

Validation and revision:
Validate the candidate.
Revise it using the diagnostics, and record changes and unresolved issues.
Preserve the acceptance criteria and initial inventory.
Submit for execution only when authorized.
If state is uncertain, do not automatically repeat an operation that might have
    already occurred.

Output:
JSON-procedure, associated worklists, explicit assumptions, validation results, and
    outstanding verification requirements.
```

### S3 Wet-lab harness for device orchestration and observation

#### S3.1 Device inventory and unified invocation

The wet-lab harness connects instruments through three control modes: API calls, preset scripts, and worklists. Device adapters translate the logical actions in a JSON-procedure into the control mode used by each instrument. The unified interface provides command invocation, execution-status queries, and result retrieval across these modes (Supplementary Fig. S2). Supplementary Table S3 lists the configured execution objects, their control modes, and the corresponding operations.

**Supplementary Table S3.** Device configuration and control modes.

| Device ID | Device or execution object | Manufacturer and model | Control mode | Invocation |
|---|---|---|---|---|
| `cytomat` | Automated incubator | Thermo Fisher Scientific; Cytomat 2C450 Lin ToS | API | Plate presentation/retraction and parameterized incubation. |
| `centrifuge` | Refrigerated centrifuge | Hettich; ROTINA 380 R ROBOTIC | API | Plate-position selection and centrifugation. |
| `pcr` | Automated thermal cycler | EASTWIN; eATC-U, 96-well module | Preset script | Registered temperature programs. |
| `kingfisher` | Nucleic-acid extraction instrument | Thermo Fisher Scientific; KingFisher Flex | Preset script | Registered extraction scripts. |
| `liquid_handler` | Liquid-handling workstation | Tecan; Freedom EVO 150 | Worklist | Deck-layout selection and run_worklist. |
| `washer` | Microplate washer | Tecan; HydroSpeed | Preset script | Named wash programs selected through a discrete parameter template. |
| `qpix` | Colony-picking system | Molecular Devices; QPix 420 | Preset script | Registered colony-picking scripts. |
| `infinite` | Microplate reader | Tecan; Infinite E Plex | Preset script | Registered optical-measurement presets and result retrieval. |
| `shaker` | Temperature-control and shaking module | EASTWIN; eMC-2000 | API | Parameterized shaking. |
| `robotic_arm` | External robotic arm | MEGAROBO; Misa-F100-6080-R2 | API | Positioning and transfer motions. |
| `rail_a` | Linear rail | MEGAROBO; MRZ-Rail4500 | API | Rail positioning and status queries. |
| `gripper_a` | Vertical gripper | MEGAROBO; Auflo-VPFinger | API | Attachment, gripping, release, and return. |
| `carousel_a` | Plate carousel | MEGAROBO; Mega-FCR200 | API | rotate_to_interaction. |
| `carousel_tips` | Tip-box carousel | MEGAROBO; Mega-FCR200 | API | rotate_to_interaction. |
| `lid_a` | Microplate lid handler | MEGAROBO; MRZ-LH04 | API | pick_up_lid and place_lid. |
| `human` | Manual-handoff notifier | Software execution object | Software handoff | send_email with handoff information. |

*Note.* Control mode identifies how the local adapter invokes experimental operations. Status queries and result retrieval are provided through the unified interface. The human entry is a software notification object.

**API control.** The harness supplies registered parameters directly to device actions. This mode supports incubation, centrifugation, shaking, and plate handling, including robotic-arm motion, rail positioning, gripping, carousel rotation, and lid handling. Device adapters expose the available actions and their parameter constraints for procedure generation and validation.

**Preset scripts.** The harness selects and invokes programs configured in the instrument-control software. Thermal-cycler programs, nucleic-acid extraction scripts, colony-picking scripts, microplate-washing programs, and reader presets are referenced by their registered names. Washer program names are assembled from the permitted combinations in a discrete parameter template; the selected name identifies the corresponding operating settings.

**Worklists.** For the Tecan Freedom EVO 150, the Dry-Lab Harness generates an Advanced GWL worklist using the selected deck layout, registered labware, Liquid Class entries, and inventory. The JSON-procedure supplies the worklist path for execution and declares dependencies between workstation operations and tasks on other devices. Validation checks the procedure and worklist together against the local configuration.

Each execution object has a unique device_id. The default configuration contains 16 objects, including separate plate and tip-box carousels and a software notifier for manual handoffs. Internal EVO 150 temperature-control and shaking modules are included in the workstation entry. The get_info, get_action_info, get_layout_info, and get_constraints calls provide capability and configuration information for planning. Supplementary Fig. S3 shows the physical platform layout.

#### S3.2 Spatial layout and labware state

Virtual Lab provides device layouts, plate-slot topology, access permissions, and transfer points. These definitions map logical identifiers to physical positions. Lab Device supplies invocation, execution-status queries, and completion records. The orchestration layer combines these interfaces to check task dependencies, device readiness, labware location, and resource availability before dispatch (Supplementary Fig. S2).

Layout definitions specify available positions and access relationships. Labware-instance records specify identity, current location, labware specification, lid status, and occupancy. Supplementary Table S4 summarizes configured logical capacities; actual occupancy is recorded in the labware state.

Confirmed transfers and material operations update runtime state and event records. Virtual Lab uses the same records for visualization and replay. Task, device, labware, slot, and execution identifiers link commands to the corresponding state changes. After an interruption, confirmed changes are retained and unresolved states are marked uncertain pending reconciliation. Manual email notifications request a handoff; loading completion and inventory changes require subsequent state confirmation.

#### S3.3 JSON procedure format and task orchestration

A JSON-procedure declares a schema version, procedure identifier, initial labware, and task collection. Optional fields provide a well-level material model, process conditions, and compatibility requirements (Supplementary Table S5). Tasks are classified as device operations, labware transfers, or wait/software-processing nodes.

The depends_on field defines the directed acyclic task graph. Tasks become eligible after their declared predecessors have succeeded, required labware is in place, devices are ready, and resources are available. Scheduling retains explicit dependencies alongside additional dependencies generated from shared process conditions.

**Supplementary Table S4.** Configured logical capacities and access constraints.

| Object or position | Configured capacity and access |
|---|---|
| Incubator | Two towers with seven storage positions each. External access is through the interaction position; plate identifiers are used to request presentation or retraction. |
| Centrifuge | Two plate positions: A for the balance plate and B for the sample plate. |
| QPix | One source-plate position and one destination-plate position. |
| Plate carousel | T01–T05: 20 positions per tower; T06–T10: 10 per tower; total 150 positions. |
| Tip-box carousel | T01–T05: 4 positions per tower; T06–T10: 6 per tower; total 50 positions. |
| Lid handler | Four sites, each maintaining the corresponding lid independently. |
| EVO 150 | The selected Table defines the layout and access constraints for LiHa, RoMa, and MCA. |
| Internal EVO 150 modules | Grid 3 / Site 1: temperature-control and shaking module. Grid 15 / Site 1: temperature-control module. Both are controlled by the EVO 150. |
| EVO 150 handoff position | Grid 6 / Site 1 supports handoff between the external robotic arm and internal RoMa. |
| Internal grids 3, 6, 8, and 15 | Internal access is restricted to RoMa. This does not remove external-arm access to the Grid 6 handoff position. |
| Grid 33 | Accessible to RoMa and MCA. |
| Other positions | Purpose and access depend on the selected Table. |

*Note.* Capacities describe configured storage positions. Labware-instance records specify occupancy.

**Supplementary Table S5.** Principal fields of the JSON-procedure.

| Field | Meaning |
|---|---|
| `schema_version` | Input-format version. |
| `id` | Procedure identifier. |
| `plates` | Labware instances and their initial locations, specifications, and lid states. |
| `tasks` | Collection of device, transfer, or wait tasks. |
| `depends_on` | Prerequisite tasks that must complete successfully. |
| `device` / `station` | Invoked device or associated station. |
| `action` / `parameters` | Registered action and its input parameters. |
| `result_action` | Interface used to query results after action completion. |
| `timeout_s` | Task timeout budget. |
| `execution_model` | Well-level model for liquids, reagents, tips, waste, and sample provenance. |
| `process_constraints` | Constraints such as cold-chain handling, mixing conditions, and well-level time windows. |

**Supplementary Note.** Illustrative interface example for plate transfer and $OD_{600}$ measurement.


```
{
  "schema_version": 1,
  "id": "transfer_and_read",
  "plates": [
    {
      "id": "P001",
      "location": "carousel_a",
      "slot": "T01-S01",
      "labware_spec": "plate_20",
      "has_lid": false
    }
  ],
  "tasks": [
    {
      "id": "select_plate",
      "kind": "device",
      "device": "carousel_a",
      "action": "rotate_to_interaction",
      "parameters": {
        "tower": 1,
        "site": 1
      }
    },
    {
      "id": "transfer_to_reader",
      "kind": "transfer",
      "plate": "P001",
      "from": "carousel_a",
      "from_slot": "T01-S01",
      "to": "infinite",
      "depends_on": ["select_plate"]
    },
    {
      "id": "read_od600",
      "kind": "device",
      "station": "infinite",
      "plate": "P001",
      "action": "read_od_600",
      "parameters": {},
      "result_action": "get_results",
      "depends_on": ["transfer_to_reader"]
    }
  ]
}
```


The example in Supplementary Note links plate presentation, transfer, and measurement through explicit dependencies. Material accounting and scientific acceptance are specified through the corresponding model and requirement fields for a complete experiment.

#### S3.4 Validation and software rule checking

Validation checks the procedure, resource state, device feedback, and acceptance criteria at submission, scheduling, execution, and completion. The interface reference snapshot contains 291 registered diagnostic-rule identifiers and 17 identifiers for independent case review or acceptance. Schema-field constraints and low-level driver safeguards supplement this catalog. A diagnostic rule can apply at several stages.

Diagnostics are grouped into eight analytical categories using their identifiers and context (Supplementary Table S6). In each evaluation and revision round, a procedure is counted once in each applicable category. Failed-procedure totals are deduplicated across categories. The 66 tasks were selected from the 219-task protocol-generation benchmark (Supplementary Fig. S4A). Supplementary Fig. S5 reports all measured rounds for the seven evaluations of the 66-task benchmark.

**Supplementary Table S6.** Diagnostic categories and principal checks.

| Category | Checks or recorded content |
|---|---|
| Format and schema | JSON syntax, duplicate keys, finite numeric values, format version, required fields, field types, and immutable context. |
| Actions and bindings | Device and action declarations, operation assets and worklist bindings, parameter combinations, and agreement between declared steps and normalized platform operations. |
| Dependencies, plate positions, and limits | DAG dependencies, required operation order, positions and occupancy, handoff inventory, and configured task limits. |
| Materials and samples | Well-level volumes and composition, sample identity and product lineage, tip state, and pipetting restrictions when labware has lids or seals. |
| Conditions and timing | Temperature, incubation and shaking conditions, waits and hold stages, sealing, cold-chain handling, and time windows across steps. |
| Analysis, units, and QC | Analysis-task execution, input–output associations, units, sample and replicate grouping, concentration, QC, and ordering constraints involving analysis tasks. |
| Other validation and coverage | Required-step coverage, pauses and handoffs, and execution, scheduling, worklist, or material diagnostics not assigned to the preceding categories. |
| Infrastructure | Executor-boundary exceptions, evaluation wall-clock timeouts, and missing execution reports expected from the infrastructure. |

*Note.* Categories summarize diagnostic content rather than sequential validation stages. Each procedure is counted at most once per category within each evaluation and revision round but may contribute to multiple categories. Total failed-procedure counts are deduplicated across categories.

Device interlocks encode dependencies between actions. For centrifugation, the balance plate is loaded into position A before the sample plate enters position B. Both plates are required for centrifugation; unloading removes the sample plate before the balance plate. The associated chain reserves the vertical gripper and returns it after use. These relationships are checked before submission or during execution.

Shared-condition scheduling accounts for the incubator temperature and $CO_2$ requirements of concurrent tasks. Matching conditions permit additional plates to enter; shaking-speed differences alone are permitted by this scheduling rule. Conflicting or unspecified shared conditions cause the later incubation to wait for the preceding task. These generated dependencies operate alongside declared depends_on constraints.

Diagnostics have ERROR, WARNING, or INFO severity. ERROR blocks validation in both normal and strict modes. Normal mode permits WARNING diagnostics, and strict mode blocks them. Catalog severities provide defaults that can be adjusted to the run context in the returned diagnostic.

Validation coverage is determined by the supplied models and requirements. The execution_model field supplies well-level material accounting. Process constraints supply cold-chain, sealing, mixing, and well-level timing requirements. Declared experimental requirements or independent case objectives define step-coverage checks. The harness reports schema validity, pre-submission validation, virtual-execution status, and independent objective acceptance as separate outputs.

##### S3.5 Completion evidence and procedure revision

The execution layer verifies action completion using the execution_id, device status, progress, and completion records. Data-producing operations also require result records associated with the current execution. Verified completion releases dependent tasks for scheduling and updates labware location, resource occupancy, and material state.

Runtime monitoring tracks timeouts, lost state updates, failures, and conflicting feedback. Unresolved completion states keep dependent tasks pending. Execution records retain task identifiers, execution identifiers, timestamps, and confirmed step-level changes, providing the basis for state display, replay, and interruption recovery.

A workflow handoff begins after the driver confirms a safe pause. Resumption checks the execution identifier, pause position, labware identity, return position, and tool state before continuing the original execution. The pause and completion states are recorded separately.

Structured diagnostics identify the rule, severity, stage, task, field path, and message. Additional fields can provide expected and observed states, repair inputs, or suggested corrections. Candidate revision uses these diagnostics to update actions, parameters, dependencies, labware bindings, or handoffs while preserving the experimental objective, confirmed inputs, and acceptance criteria. Additional information and state-confirmation requirements remain recorded until resolved.

Before execution, the harness checks the validated procedure against the current configurations, files, and state. After interruption, recovery reconciles existing execution records with the current platform state, retains confirmed changes, and resolves uncertain operations before any repeat execution. Candidate procedures, diagnostics, revisions, and run records retain the complete generate–validate–revise history.

### S4 Protocol evaluation on the 219-task benchmark

The benchmark contains 219 tasks assigned to eight primary experimental domains (Supplementary Fig. S4A). Reference protocols were prepared and reviewed for the local devices, reagents, and consumables, then converted into experimental intents. Three generation-and-review rounds used the same task set, with Review agent error analyses informing generation in the next round.

Three human experts and the Review agent independently scored each protocol using the five-dimensional, 100-point rubric in Supplementary Table S7. The human score $H$ was the mean of the three expert scores. The composite score $S$ combined $H$ with the agent score $A$:

$$H = \frac{H_1 + H_2 + H_3}{3}, \qquad S = \frac{2}{3}H + \frac{1}{3}A. \tag{1}$$

A protocol passed when the human and agent assessments both approved it, the composite score reached 80/100, and the review confirmed that essential experimental requirements were satisfied. A confirmed critical issue triggered rejection regardless of the score. Critical issues were substantiated errors or omissions that could prevent an essential operation, invalidate a key result, or compromise interpretation. Quality-control requirements were evaluated in the context of the task (Supplementary Table S8).

**Supplementary Table S7.** Protocol evaluation criteria and maximum scores.

| Evaluation dimension | Maximum points | Assessment focus |
|---|---|---|
| Requirement consistency | 30 | Adherence to the experimental objective, stated requirements, and available local devices, reagents, and consumables. |
| Scientific rationality | 25 | Scientific soundness of selected methods, experimental conditions, parameter choices, and quantitative reasoning. |
| Completeness and executability | 25 | Completeness and logical order of essential steps, required inputs and quantities, and feasibility of the resulting procedure. |
| Quality control and exception handling | 10 | Appropriate controls and quality checks, with interpretation criteria and responses to outcomes where relevant to the task. |
| Clarity and traceability | 10 | Clear units and sample and reagent identities, internally consistent instructions, and traceability of the proposed workflow. |
| Total | 100 | |

**Supplementary Table S8.** Representative critical issues across the three protocol-review rounds.

| Review round | Experimental task | Reviewer-identified issue and assessed consequence |
|---|---|---|
| 1 | Golden Gate assembly | Restriction-site design: a BsaI recognition site remained in the target-vector backbone, creating a risk of re-cleavage during assembly. |
| 1 | GST pull-down | Sample-volume planning: only 20 $\mu$L of eluate was available, but two 20 $\mu$L sample loads were planned; the available material was insufficient. |
| 1 | CUT&Tag | Biological replication: cells were pooled before being split into nominal biological replicates, preventing assessment of independent biological variation. |
| 2 | Golden Gate assembly | Primer structure: the required spacer between the BsaI recognition site and the 4-bp overhang was omitted, compromising the intended cleavage-product design. |
| 2 | CUT&Tag | QC consumption omitted from material accounting: removing 10 $\mu$L from a 55 $\mu$L reaction left 45 $\mu$L, which was insufficient for the next planned step. |
| 3 | GST pull-down | Essential detection steps omitted: the procedure proceeded from elution directly to membrane blocking without electrophoresis and transfer, leaving the Western blot workflow incomplete. |
| 3 | AST activity assay | Insufficient sample: preliminary testing required 144 $\mu$L of serum, whereas only 120 $\mu$L was available. |
| 3 | Enzyme activity assay | Inconsistent volume units: some reagent quantities lacked units; interpreting them as $\mu$L would introduce a 1,000-fold discrepancy. |
| 3 | Wound-closure assay | Unspecified QC assessment: a cell-viability threshold of at least 90% was required without a measurement method or calculation rule. |

### S5 Biological workflows and supporting data

#### S5.1 Automated workflows and execution timelines

Supplementary Fig. S6 shows the automated workflows and representative execution timelines.

#### S5.2 Vector development, sample preparation, and screening

Supplementary Fig. S7 provides supporting data for vector development, clone selection, titer measurement, normalization, and ELISA screening.

#### S5.3 Titer-guided normalization strategy

Supplementary Fig. S7G shows the normalization scheme used before ELISA. The scheduling comparison and cumulative changes in v0–v4 are described in the main-text section on phage titer normalization scheduling. These comparisons use one run per version in each of the three layouts (Original, Layout 1, and Layout 2) and report summed action times rather than total elapsed runtime.
